# Isogenic forebrain organoids uncover early neurodevelopmental alterations and imbalances in neuronal function leading to hyperexcitation in Gaucher disease

**DOI:** 10.64898/2026.09.27.754744

**Authors:** James A. Crowe, Zuzana Matusova, Aline S. Girard, Amélie Rajon, David Colameo, Leal Oburoglu, Oskar G. Zetterdahl, Joachim Björklund, Claudia Puigsallosas Pastor, Svenja Krupp, Ulf Ellervik, Alessio Cremonesi, Daniel Tornero, David Penton Ribas, Henrik Ahlenius, Lukas Valihrach, Isaac Canals

**Affiliations:** Division of Metabolism, University Children’s Hospital Zurich, University of Zurich, Zurich, Switzerland; Zurich Neuroscience Center, Zurich, Switzerland; Department of Experimental Medical Sciences, Lund University, Lund, Sweden; Institute of Biotechnology of the Czech Academy of Sciences, Vestec, Czech Republic; Electrophysiology Facility (e-phac), University of Zurich, Zurich, Switzerland; Institute for Regenerative Medicine, University of Zurich, Zurich, Switzerland; Department of Chemistry, Lund University, Lund, Sweden; Laboratory of Neural Stem Cells and Brain Damage, Department of Biomedical Sciences, Institute of Neursciences, University of Barcelona, Barcelona, Spain; Institut d’Investigacions Biomèdiques August Pi i Sunyer (IDIBAPS), Barcelona, Spain; Centro de Investigación Biomédica en Red de Enfermedades Neurodegenerativas, Instituto de Salud Carlos III, Madrid, Spain; Division of Clinical Chemistry and Biochemistry, University Children’s Hospital Zurich, University of Zurich, Zurich, Switzerland

**Author notes:** Department of Molecular Life Sciences, URPP AdaBP, University of Zurich, Zurich, Switzerland.

## Abstract

Gaucher disease is a rare lysosomal storage disorder caused by autosomal recessive mutations in the *GBA1* gene, encoding the lysosomal enzyme glucocerebrosidase. Gaucher disease is classified in 3 different subtypes depending on the presence and severity of neurological involvement, with type 2 resulting in fatal early-onset neuropathology and patients exhibiting developmental delays, seizures and early death. Studies investigating disease mechanisms of neuronopathic Gaucher disease are mainly based on animal models and focus predominantly on late neuronal phenotypes. Here, we established healthy control and Gaucher disease patient-derived iPSC lines and engineered them to obtain isogenic control and disease lines. Using these lines, we generated cortical and subpallial brain organoids in which we identified early-onset lipid dysregulation in form of glucosylceramide accumulation, highly elevated glucosylsphingosine, and a later increase in ganglioside levels, recapitulating clinical findings. Furthermore, single-cell transcriptomic profiling uncovered novel phenotypes in both cortical and subpallial forebrain organoids. Subpallial alterations consisted of an early increase in migrating interneurons in subpallial organoids, which upregulated cholesterol metabolism. Cortical alterations showed early upregulation of mitochondrial genes and a downregulation of proliferation, with a subsequent switch from GABAergic to glutamatergic neuron fate with a striking increase in gene expression related to the synaptic assembly. Functional assays demonstrated a marked hyperexcitability of cortical organoids and reduced response to GABA-A receptor blockage in Gaucher disease. Additional 2D neuronal network models confirmed the organoid data and showed that both glutamatergic and GABAergic neurons contribute to the phenotype, with hyperexcitability of Gaucher glutamatergic neurons and incapacity of Gaucher GABAergic neurons to balance the excessive excitation. This alteration represents a clinically significant phenotype as many patients exhibit an excitation/inhibition imbalance leading to treatment-resistant seizures, hastening their decline. In conclusion, our defined human models of Gaucher disease identify novel and clear phenotypes that can be used for drug screening or aid in development of new therapeutic strategies to ameliorate Gaucher disease.

## INTRODUCTION

Lysosomal storage disorders (LSDs) encompasses a group of rare inherited metabolic diseases caused by deficiencies in lysosomal enzymes, leading to accumulation of undegraded molecules and impairment of the lysosomal system^1^, which is essential for many cellular processes^2^. Gaucher disease is the most common lysosomal storage disorder caused by mutations in the *GBA1* gene, which encodes the enzyme β-glucocerebrosidase (GCase)^3^. In addition, carrying *GBA1* mutations has been linked to a higher risk of developing Parkinson’s disease^4^. Deficiency of GCase results in the accumulation of glucosylceramide (GlcCer) within lysosomes, ultimately disrupting normal cellular function and triggering a range of symptoms such as hepatosplenomegaly, pancytopenia and bone alterations^5, 6^. Gaucher disease is classified into three main types: type I (non-neuronopathic) Gaucher disease is associated primarily with anemia, bone disease, and organ enlargement, but spares the nervous system; type II (acute neuropathic) is the most severe form, affecting mostly infants with rapid neurodegeneration and causing very early mortality; and type III (chronic neuropathic) with a more variable course, later onset and slower progression of neurological symptoms. Currently, treatment options include enzyme replacement and substrate reduction therapies, which can significantly improve systemic symptoms but have little effect on neurological manifestations^7^. This is because enzymes cannot effectively cross the blood-brain-barrier, leaving the brain untreated and vulnerable to disease progression in patients of type II and type III Gaucher disease, and presenting a putative explanation of the increased risk of Parkinsonian symptoms seen in type I Gaucher disease patients^8^. The lack of effective brain-targeted therapies highlights the urgent need for better disease models that accurately replicate the neurological aspects of Gaucher disease and assist in developing and testing new treatment strategies.

GlcCer is a glycosphingolipid formed by the enzyme glucosylceramide synthase, which adds a glucose residue to ceramide^9^, and it is found mainly in the Golgi and in the cell membranes. Within the Golgi, GlcCer formation is the first step in the synthesis of complex glycosphingolipids such as gangliosides^9^, which are molecules essential in key pathways for normal brain development and functionality of brain cells^10^. Degradation of GlcCer occurs within lysosomes, and when impaired, GlcCer accumulates and can be deacetylated, generating glucosylsphingosine (GlcSph), which has been suggested to be neurotoxic and it is a biomarker of Gaucher disease^11^.

For many years, studies on disease mechanisms of Gaucher disease have used animal models, specifically mice^12^, although mainly focused on neuronal phenotypes after the onset of the disease. However, one study identified a decrease in brain-derived neurotrophic factor (BDNF) and nerve growth factor (NGF) during brain development in mice^13^, which could contribute to neuronal loss in neuronopathic Gaucher disease patients. Despite their valuable insights, animal models do not fully recapitulate the human disease timeline and neuronopathic phenotypes^12^, possibly due to species-specific differences as well as the reduced cellular and organ complexity when comparing the human and murine brain^14–18^.

A major obstacle to study neurological phenotypes of Gaucher disease is the difficulty to obtain human brain tissue for research. The use of human iPSCs has allowed scientists to model several disorders *in vitro* using protocols to differentiate iPSCs towards mixed neural cell types by mimicking developmental cues. In addition, and to better recapitulate the spatial organization, cellular diversity, and cell-cell interactions of the developing human brain, protocols to generate brain organoids from iPSCs have been developed^19^. To better understand the neuropathology of human Gaucher disease, several human iPSC-derived models have been generated, including neurons^20–28^ and astrocytes^29^. Moreover, a few studies have also generated midbrain organoids from iPSC lines carrying heterozygous *GBA1* mutations for investigating Parkinson’s disease mechanisms^30–32^, and recently, also Gaucher disease^33^. In this last study, authors showed that midbrain organoids displayed lipid accumulation and transcriptomic changes, with subsequent impairments in dopaminergic neuron differentiation. However, to our knowledge, there are no studies using Gaucher iPSC lines to generate organoids representative of other brain regions for identifying early neurodevelopmental alterations arising from dysregulated glycosphingolipid metabolism.

Here, we successfully generated iPSCs from fibroblasts of a Gaucher disease patient. We used genome editing to correct the disease causative mutations in the patient line and to knockout the *GBA1* gene in a healthy control line we previously established, thus generating isogenic control and disease lines respectively. We have differentiated these four iPSC lines towards cortical and subpallial forebrain organoids and showed that they recapitulate some of the hallmarks of the disease such as a clear accumulation of GlcSph with a subsequent increase in gangliosides, specifically GM_1_. These changes in lipids were followed by alterations in neural lineage specification, pathways related to cell migration, proliferation, maturation and functionality of neurons, which were identified after single-cell transcriptomics. Finally, functional assays demonstrated important functional deficits in both glutamatergic and GABAergic neurons in neuronopathic Gaucher disease, providing new knowledge on disease mechanisms and cell-type specific contributions.

## RESULTS

### Generation of healthy control and Gaucher disease iPSC lines with corresponding isogenic controls

To generate novel human models of neuronopathic Gaucher disease, we first reprogrammed one disease (GD) skin fibroblast line into iPSCs using mRNA reprogramming to be used together with our previously generated healthy control iPSC line^34^. Clonal lines were established, expanded, and fully characterized following the International Society for Stem Cell Research (ISSCR) guidelines^35^. Colonies displayed typical stem cell morphology (Fig S1A), expression of markers of undifferentiated state NANOG, OCT3/4, SOX2 and TRA-1-81 at the protein level (Fig S1A), and SOX2, NANOG and POU5F1 at the mRNA level (Fig S1B). The GD line did not show any chromosomal abnormalities (Fig S1C) and had the capacity to differentiate towards the three germ layers (Fig S1D). Sanger sequencing of the *GBA1* gene confirmed the presence of the patient mutations, P415R and L483P (historically known as P454R and L444P, respectively) (Fig S1E). Surprisingly, after Sanger sequencing the *GBA1* gene in the healthy control line, we discovered that this individual is a carrier for one of the most frequent mutations, N370S (historically known as N370S) (Fig S1F), so we designated this line as a healthy carrier (HC).

We next used CRISPR/Cas9 genome editing to generate an isogenic KO line by disrupting the *GBA1* gene in the HC line. The presence of a pseudogene (*GBA1LP*) with 96% of homology to the *GBA1* gene 16 kb downstream, represents a challenge to specifically target the gene. We designed a strategy targeting a 55-bp region in *GBA1* exon 9 that is missing in the *GBA1LP* to disrupt the *GBA1* gene in the HC line (Fig 1A), identifying one clone in which both alleles had the *GBA1* gene disrupted. One allele had a 74-bp insertion leading to a premature stop codon (K447fsX21) while the other allele had a 30-bp deletion leading to the loss of 10 amino acids and the substitution of another one (F436_T446delinsS, Fig 1B). We designated this isogenic disease-like line as HC-KO.

**Figure 1.**
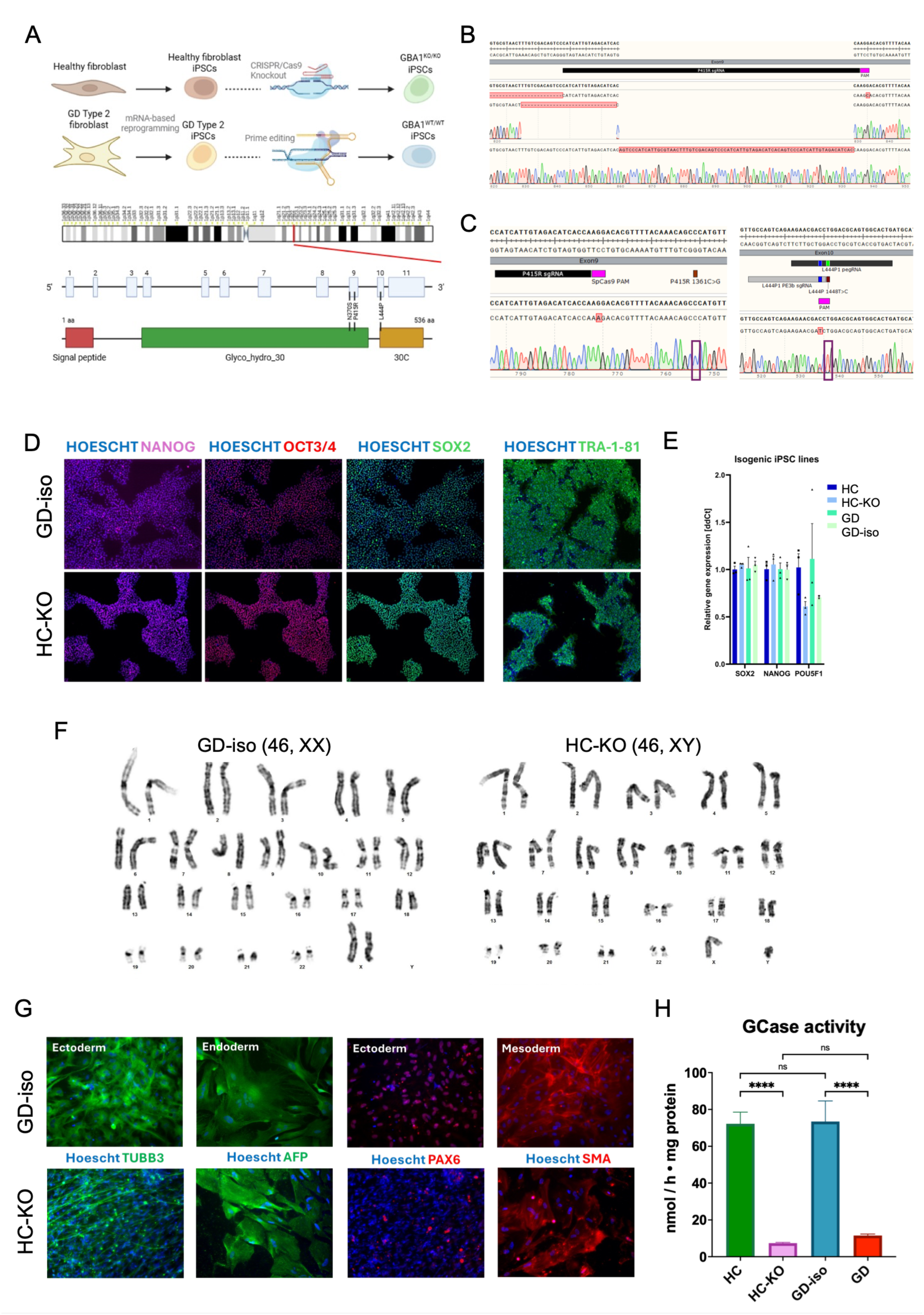
Generation of isogenic lines with CRISPR/Cas9 genome editing. **A)** Schematic representation of the CRISPR strategy to knock out the GBA1 gene in the healthy control iPSC line and to correct the mutations in the GD iPSC line (upper image); and schematic representation of chromosome 1 with a zoom-in at the region where the GBA1 gene is located with the patient mutations highlighted (P415R and L444P). **B)** Chromatogram showing the sequencing of the HC-KO line with the two alleles and the corresponding deletion in each one. **C)** Chromatogram showing the correction of the mutations in the GD-iso line. **D)** Representative images of immunocytochemistry for the markers of undifferentiated state NANOG, OCT3/4, SOX2 and TRA-1-81 of the iPSC for the GD-iso and HC-KO lines. **E)** Relative expression of the markers of undifferentiated state SOX2, NANOG and POU5F1 normalized to GAPDH. **F)** Representative image of the G-band karyotyping for the GD-iso and HC-KO iPSC lines. **G)** Representative images of immunocytochemistry for markers of ectoderm (TUBB3, PAX6), endoderm (AFP) and mesoderm (SMA) after trilineage differentiation for both GD-iso and HC-KO lines. **H)** Enzymatic assay for GCase activity after 7 days of differentiation of all lines towards neurons showing nmol of product per h and mg of protein. All bar graphs represent the mean ± s.e.m of 3 independent experiments. **** P<0.0001, one-way ANOVA comparing all groups.

The 55-bp region specific to the *GBA1* gene is in very close proximity (41 bp upstream) to the P415R mutation of the GD line (Fig 1A), thus the same editing strategy was used to correct this mutation (Fig 1C). To correct the second mutation (L444P), located within exon 10 where there are no mismatches in the *GBA1* compared to the *GBA1LP*, we chose to used prime editing (PE3b) for its specificity, to avoid double strand breaks, and for the low risk of off-target effects^36^. We identified one clone in which the second mutation, L444P, had been corrected (Fig 1C). We named this isogenic control line GD-iso.

After generating the HC-KO and the GD-iso lines, we confirmed expression of markers of undifferentiated state both at the protein (Fig 1D) and mRNA (Fig 1E) levels, lack of chromosomal abnormalities (Fig 1F), and capacity to differentiate towards the three germ layers (Fig 1G) for both lines.

To validate the isogenic lines, we performed a GCase enzymatic activity assay (Fig 1H), showing that the HC line had an enzymatic activity of 72.23 nmol / h x mg protein, much higher than the activity of the GD line (11.57 nmol / h x mg protein). HC-KO derived cells displayed very low GCase activity (7.39 nmol / h x mg protein), while the GD-iso enzymatic activity was restored to control levels (73.46 nmol / h x mg protein).

Altogether, using mRNA reprogramming and genome editing, we successfully generated 2 pairs of healthy-disease isogenic iPSC lines (Table S1), which constitute a valuable resource to investigate neuronopathic Gaucher disease mechanisms.

### Gaucher disease brain organoids present progressive sphingolipid accumulation

To generate novel models of neuronopathic Gaucher disease, we differentiated all iPSC lines (HC, HC-KO, GD and GD-iso) towards human forebrain subpallial (hSOs) and cortical (hCOs) organoids (Fig 2A) using a previously established protocols^37^. We maintained hSOs and hCOs for up to 150 days in vitro (DIV), with both hSOs and hCOs growing similarly between control and patient lines (Fig 2B). All lines generated SOX2+ neural rosettes and NeuN+ neurons at DIV75 (Fig 2C,D).

**Figure 2.**
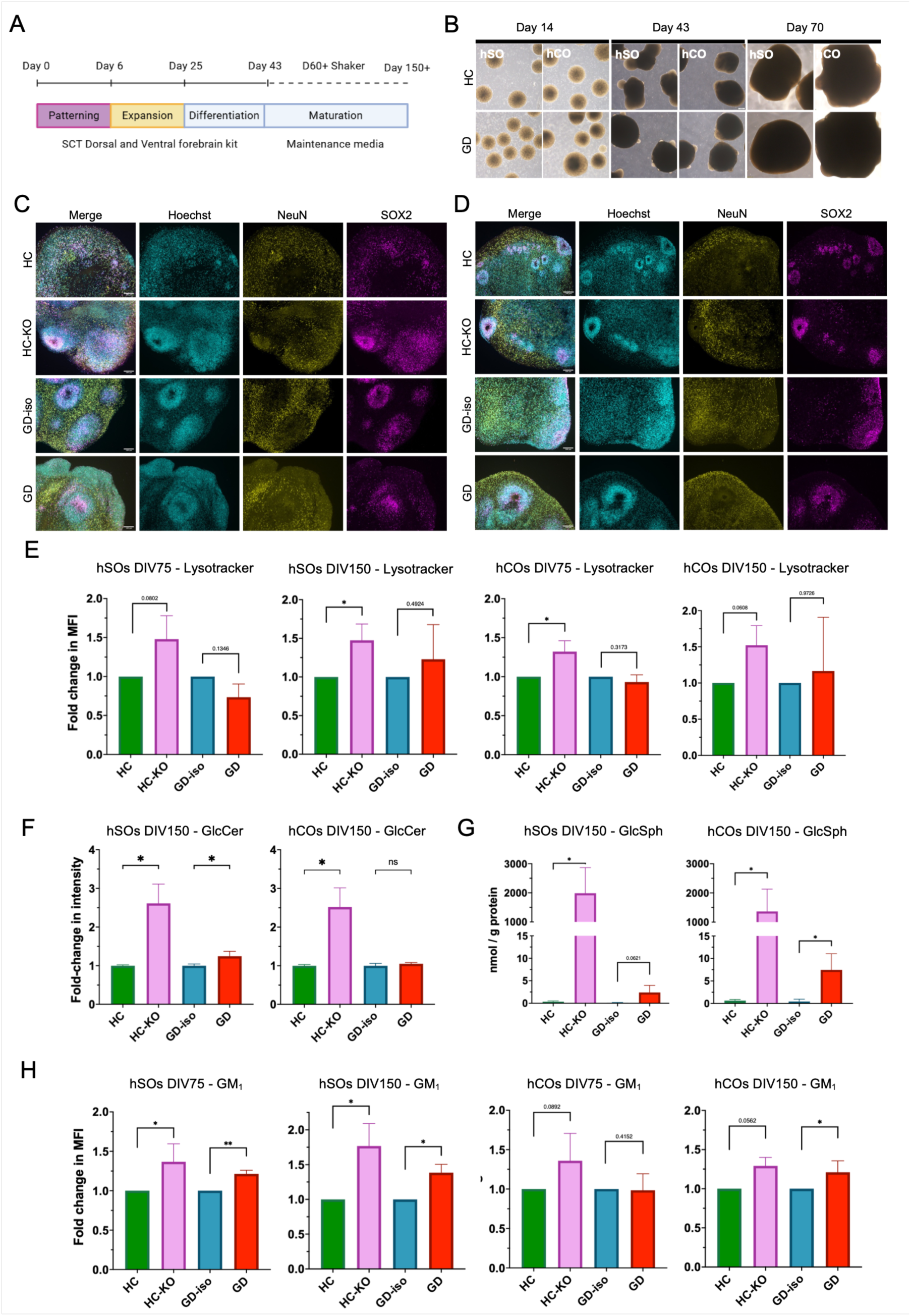
Gaucher disease forebrain organoids display progressive lipid accumulation. **A)** Schematic representation of the generation of hSO and hCO. **B)** Representative brightfield images at different days of differentiation of hSO and hCO for the HC and the GD lines. **C)** Representative images of immunocytochemistry for Hoechst (nuclei), NeuN (neurons) and SOX2 (neural stem cells) for hSO generated from all lines at DIV75. Scale bar = 100 µm. **D)** Representative images of immunocytochemistry for Hoechst (nuclei), NeuN (neurons) and SOX2 (neural stem cells) for hCO generated from all lines at DIV75. Scale bar = 100 µm. **E)** Fold change in fluorescent intensity of Lysotracker staining in hSOs and hCOs generated from all lines at DIV75 and DIV150 comparing HC-KO to HC and GD to GD-iso. Bar graphs represent the mean ± s.e.m. of 3 independent experiments. *p<0.05, one-tailed ratio paired t-test. **F)** Fold change in dot blot intensity for GlcCer in hSO and hCO at DIV150 comparing HC-KO to HC and GD to GD-iso. Bar graphs represent the mean ± s.e.m. of 3 independent experiments. *p<0.05, one-tailed ratio paired t-test. **G)** GlcSph quantification in nmol per g of protein in hSO and hCO at DIV150 comparing HC-KO to HC and GD to GD-iso. Bar graphs represent the mean ± s.e.m. of 3 independent experiments. *p<0.05, one-tailed ratio paired t-test. **H)** Fold change in intensity of GM_1_ lipid raft staining in hSO and hCO at DIV150 comparing HC-KO to HC and GD to GD-iso. Bar graphs represent the mean ± s.e.m. of 3 independent experiments. *p<0.05, one-tailed ratio paired t-test.

An increase in the lysosomal size and number is a hallmark of many lysosomal storage disorders including Gaucher disease. To investigate this, we analysed the lysosomal content of organoids at early (DIV75) and later stages (DIV150) of organoid development by FACS using Lysotracker dye. We observed that at DIV75 and DIV150 for both types of organoids (hSOs and hCOs), only the HC-KO line presented a tendency or a significant increase in Lysotracker signal compared to its isogenic line with a 1.48-fold increase in hSO-DIV75, 1.32-fold increase in hCO-DIV75, 1.48-fold increase in hSO-DIV150 and 1.52-fold increase in hCO-DIV150 (Fig 2E). Meanwhile, for the GD and GD-iso organoids, there were no clear differences in Lysotracker signal. This might be explained by the KO having a more severe phenotype due to lower enzymatic activity as compared to GD (Fig 1H).

Gaucher disease is caused by a deficiency in GlcCer catabolism within the lysosome, leading to its accumulation. We performed a dot blot analysis to assess GlcCer accumulation in hSOs and hCOs at DIV150 when there is a clear lysosomal increase (Fig 2F, Fig S2A), and found a striking increase in GlcCer levels for HC-KO compared to HC (2.6- and 2.5-fold increase in hSO and hCO, respectively) but more modest or unclear increase in GD organoids compared to GD-iso (1.25- and 1.05-fold increase in hSO and hCO, respectively). In Gaucher disease there is also accumulation of GlcSph, the deacetylated form of GlcCer, which is used as a biomarker^11^. We quantified GlcSph using HPLC-MS at DIV150 (Fig 2G), detecting significantly high levels in both GD (2.4 and 7.46 nmol / g of protein in hSO and hCO, respectively) and HC-KO (1987.75 and 1364.55 nmol / g of protein) compared to GD-iso (0.08 and 0.45 nmol / g of protein) and HC organoids (0.34 and 0.64 nmol / g of protein). Interestingly, for both GlcCer and GlcSph, HC-KO presented a much higher degree of accumulation, again suggesting a more severe phenotype in HC-KO as compared to GD. For GlcSph, differences between GD and GD-iso were already detected at DIV75 (Fig S2B), indicating that GlcSph is already accumulated during early brain development.

These results prompted us to investigate metabolic pathways related to GlcCer, focusing on gangliosides, complex lipids that are very abundant in the brain, for which GlcCer is the biosynthetic precursor^10, 38, 39^, and that accumulate secondarily in Gaucher disease^6^. We first performed thin layer chromatography of whole-cell lysates and detected several ganglioside forms, with GM_1_ being the most abundant. Our data indicated a possible increase of GM_1_ in disease hSOs and hCOs, mainly at DIV150 (Fig S2C). To further examine the fraction of GM_1_ at the cell membranes, where gangliosides are enriched within lipid rafts and are essential for supporting, axonal growth, cell migration, synaptogenesis, and neurotransmission, we performed FACS assays using Vybrant Alexa Fluor 555 Lipidraft labelling. We detected a clear increase in fluorescence in hSOs at DIV75 for both disease lines (1.37-fold increase in HC-KO vs. HC and 1.21-fold increase in GD vs. GD-iso), and an increase in fluorescence in both hSOs (1.77-fold increase in HC-KO vs. HC and 1.39-fold increase in GD vs. GD-iso) and hCOs (1.29-fold increase in HC-KO vs. HC and 1.21-fold increase in GD vs. GD-iso) at DIV150 (Fig 2H). These results indicate that GM_1_ progressively accumulates in the membranes of cells within both types of Gaucher disease brain organoids due to the dysfunction in GlcCer catabolism.

Altogether, our data indicates an early accumulation of disease biomarkers and a progressive increase of complex sphingolipids upstream of GlcCer in brain organoid models of Gaucher disease. These changes occur before the onset of a clear lysosomal accumulation and are more striking in a KO line than a patient-derived line, as well as a seemingly earlier onset in subpallial versus cortical brain regions.

### Gaucher disease hSOs present increased number of migrating interneurons

Glycosphingolipids such as gangliosides and GlcCer are essential for normal brain development. To examine cell type identities and lineage commitment within our hSOs and hCOs and how these are affected in Gaucher disease, we next performed single-cell RNA sequencing analysis of DIV75 and DIV150 organoids from 3 independent differentiations using the 4 iPSC lines to minimize batch-to-batch and line-to-line variability. To facilitate the processing of samples from independent differentiations and different days in vitro, we used the 10x Fixed RNA Profiling system, allowing to fix samples at different days but to process them simultaneously. Thus, we extracted data from 21295 Gaucher and 17939 control cells for hSOs and from 20322 Gaucher and 23017 control cells for hCOs.

The sequencing data identified 9 cell populations in hSOs, including cycling cells, stressed cells, radial glia, astroglia, ventral progenitors, migrating interneurons, cholinergic neurons, striatal MSN neurons and GABAergic neurons (Fig 3A-B). All clusters were annotated using cell-type specific markers (Fig 3D). Both control and Gaucher hSOs showed an increase in the number of ventral progenitors and migrating interneurons over time, from DIV75 to DIV150 (Fig S3A). Interestingly, comparing Gaucher disease and control hSOs, at DIV75 there was a decrease in striatal MSN neurons and a marked increase in migrating interneurons (Fig 3C and Fig S3D), while at DIV150 the only differences were a slight decrease in cholinergic neurons and an increase in striatal MSN neurons (Fig S3B), two populations present in very small numbers at that time point (Fig S3G).

**Figure 3.**
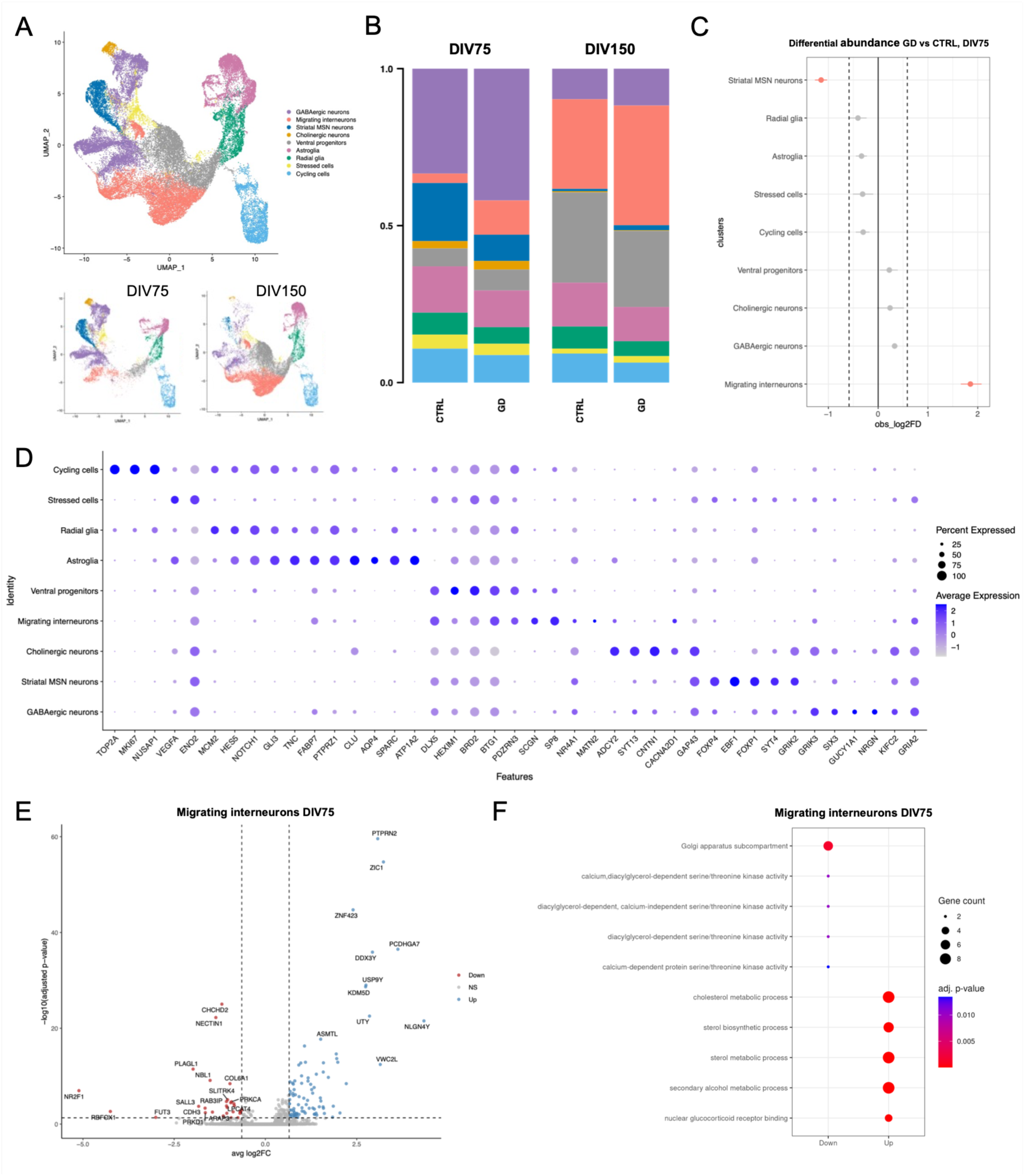
Single-cell RNA sequencing of hSO reveal changes in the number of migrating interneurons and alterations in signaling pathways and cholesterol metabolism in these cells. **A)** UMAP plot showing 9 clusters and the distribution of cells for DIV75 (lower left) and DIV150 (lower right). GD (GD + HC-KO) = 21295 cells, CTRL (GD-iso + HC) = 17939 cells. **B)** Proportional bar plots showing the abundance of cells for each cluster in each genotype group at DIV75 and DIV150. **C)** Differential abundance of populations comparing GD vs CTRL at DIV75. **D)** Dot plot with selected marker genes for each cluster. **E)** Volcano plot displaying upregulated (up, blue) and downregulated (down, red) in migrating interneurons comparing GD vs CTRL at DIV75. **F)** Overrepresentation analysis results showing top five downregulated (down) and upregulated (up) GO terms distinguishing migrating interneurons from GD vs CTRL at DIV75.

We next investigated differentially expressed genes (DEGs) at both DIV75 and DIV150 in the whole dataset, identifying 114 upregulated and 70 downregulated genes at DIV75 and 106 upregulated and 22 downregulated genes at DIV150 (Fig S3C,D,F,G). Interestingly, the top downregulated gene at both timepoints was NR2F1 (Fig S3C,F), which has been implicated in regulating the balance between excitatory and inhibitory neurons^40^. Gene ontology (GO) enrichment at DIV75 pointed towards a downregulation of plasma membrane-related processes and an upregulation of lysosomal- and synaptic-related pathways (Fig S3E). At DIV150, GO clearly indicated an upregulation of synaptic functional pathways and a downregulation of pathways related to regionalization, migration and extracellular matrix (Fig S3H).

Since the most significant population changes were found in migrating interneurons at DIV75, we decided to analyse this population at this specific timepoint in more detail. Our analysis showed 89 upregulated genes and 29 downregulated genes (Fig 3E and Fig S3D), with GO enrichment showing cholesterol metabolism pathways as the top upregulated processes together with synaptic adhesion, while the main downregulated pathways were related to Golgi and autophagy function as well as PKC signalling (Fig 3F).

Altogether, our transcriptomic analysis of forebrain subpallial organoids showed all the expected populations within these organoids and pointed towards changes in the number of migratory interneurons during brain development in Gaucher disease, with alterations in cholesterol metabolism and specific signalling pathways that play key roles for developmental processes.

### Gaucher disease hCOs display early upregulation of mitochondrial genes and a developmental switch from GABAergic to glutamatergic neuron fate

We next focus on hCOs, where data identified 14 cell populations including cycling cells, stressed progenitors, undefined progenitors, multipotent progenitors, radial glia, OPC, astroglia, early progenitors, undefined neurons, Cajal-Retzius neurons, intermediate progenitors, glutamatergic neurons, ventral progenitors and GABAergic neurons (Fig 4A). All clusters were annotated using cell-type specific markers (Fig 4D). From DIV75 to DIV150, both control and Gaucher hCOs showed a decrease in the presence of Cajal-Retzius neurons and an increase in GABAergic neurons, in line with known developmental trajectories (Fig 4B and FigS4A). Interestingly, comparing Gaucher disease and control hCOs, a high number of GABAergic neurons versus glutamatergic neurons was detected in Gaucher organoids at DIV75 while at DIV150, there was a clear increase in excitatory lineage (glutamatergic neurons and intermediate progenitors) at the expense of inhibitory lineage (ventral progenitors and GABAergic neurons) (Fig 4C, Fig S4B). These results suggest an alteration in neuronal lineage commitment throughout development resulting in changes in proportions of the different neuronal subtypes.

**Figure 4.**
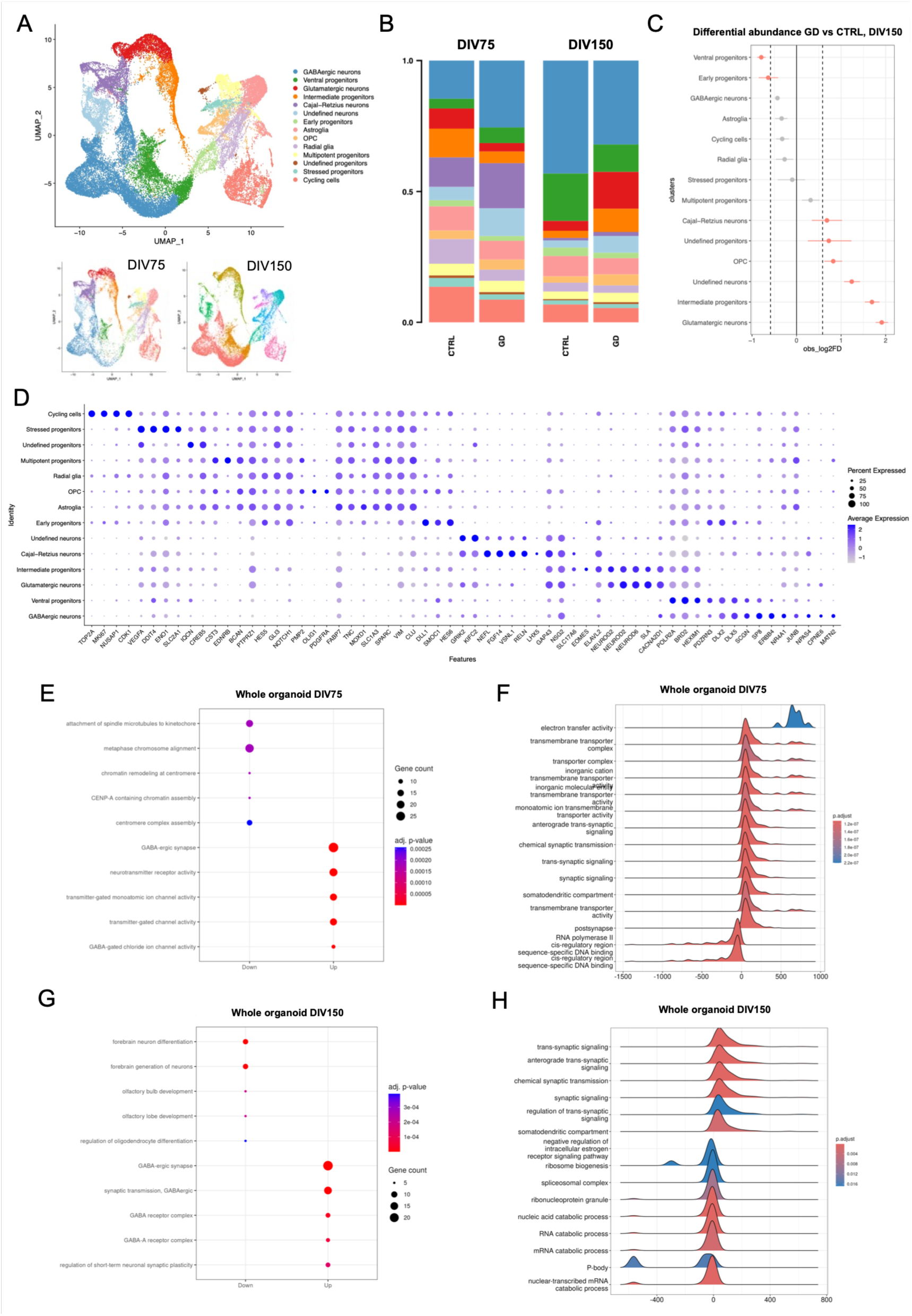
Single-cell RNA sequencing of hCO reveal changes in neuronal proportions and alterations proliferation, differentiation and maturation of neurons. **A)** UMAP plot showing 14 clusters with the distribution of cells for DIV75 (lower left) and DIV150 (lower right). GD (GD + HC-KO) = 20322 cells, CTRL (GD-iso + HC) = 23017 cells. **B)** Proportional bar plots showing the abundance of cells for each cluster in each genotype group at DIV75 and DIV150. **C)** Differential abundance of populations comparing GD vs CTRL at DIV150. **D)** Dot plot with selected marker genes for each cluster. **E)** Overrepresentation analysis results showing top five downregulated (down) and upregulated (up) GO terms distinguishing whole organoids from GD vs CTRL at DIV75. **F)** Ridge plot showing the top 15 enriched Biological Processes (GO: BP) identified by GSEA. Genes were ranked according to their log2FoldChange (log2FC). The log2FC distribution for each GO term indicates whether the biological process is mainly upregulated (positive values) or downregulated (negative values) in GD compared to CTRL at DIV75. **E)** Overrepresentation analysis results showing top five downregulated (down) and upregulated (up) GO terms distinguishing whole organoids from GD vs CTRL at DIV150. **F)** Ridge plot showing the top 15 enriched Biological Processes (GO: BP) identified by GSEA. Genes were ranked according to their log2FoldChange (log2FC). The log2FC distribution for each GO term indicates whether the biological process is mainly upregulated (positive values) or downregulated (negative values) in GD compared to CTRL at DIV150.

DEG analyses of whole organoids showed a total of 659 upregulated and 1008 downregulated genes at DIV75 and 498 upregulated and 72 downregulated genes at DIV150 (Fig S4C-F). GO enrichment of DIV75 organoids showed a downregulation of pathways related to cell cycle paired with an upregulation of synaptic genes, clearly pointing towards an accelerated differentiation within Gaucher organoids compared to control organoids (Fig 4E). In addition, gene set enrichment analysis (GSEA) demonstrated a strong upregulation of electron transfer activity (Fig 4F), in line with a previous study showing that GCase import is important in midbrain organoids to maintain mitochondrial function^27^. These differences were related to upregulation of a set of 10 closely related mitochondrial genes (Fig S4C, S5A), and the cell clusters expressing high levels of these genes were mainly Cajal-Retzius neurons and undefined neurons, which were specifically charaterized by high expression of these genes. The latter could represent highly stressed cells from the core of the organoids. However, the first group represents highly migratory cells, suggesting that the dysregulation in mitochondrial genes can have an effect in their migratory capacity in Gaucher disease as shown for migratory interneurons in hSOs. An increase in mitochondrial genes can also represent the metabolic switch from glycolisis to oxidative phosphorylation occuring during neuronal differentiation. Supporting this idea, GD organoids have a lower number of progenitors, which could suggest increased differentiation rate. To pinpoint the contribution of the two major neuronal types to these changes, we examined TMRM intensity, in 2D cultures of glutamatergic and GABAergic neurons independently, identifying an increase in TMRM staining in both types of neurons in GD (Fig S5B,C). This increase in TMRM can be due either to a higher mitochondrial membrane potential and potentially an increase in oxidative phosphorylation, or to an increase in the number of mitochondria.

GO enrichment analyses comparing Gaucher to healthy hCOs at DIV75 also showed an upregulation in pathways related to GABAergic synaptic and neurotransmission functions (Fig 4E), in line with the idea of a faster maturation of neurons and formation of neuronal networks. Meanwhile, top downregulated genes were related to cell cycle (Fig 4E), suggesting impaired proliferation capacity within Gaucher hCOs. GO enrichment analysis at DIV150 highlighted upregulation in processes related to GABAergic inhibitory neuron development (Fig 4G). These results are interesting since Gaucher disease hCOs showed fewer GABAergic neurons and an increased number of glutamatergic neurons, which suggests a potential compensation mechanism to overcome the lack of GABAergic neurons. This was exemplified by a downregulation of key GABAergic lineage genes (*DLX1/2/5, SLAIN1, SP9, TOX3*), and upregulation of glutamatergic lineage genes (*BHLHE22, EMX1, NEUROD2/6, SATB2*). In contrast, downregulated GO terms were related to brain development and differentiation (Fig 4G), indicating a potential acceleration of maturation in line with the mitochondrial phenotype at DIV75. GSEA analysis supported these results by showing an enrichment in pathways related to synapse formation and function and under-representation of pathways related to cell cycle (Fig 4H), again supporting the accelerated differentiation phenotype.

Altogether, the results from single-cell RNA sequencing experiments clearly indicate an early alteration in mitochondria in Gaucher hCOs together with decreased proliferation and increase differentiation/maturation, followed by a later loss of GABAergic neurons together with an increase in glutamatergic neurons and an increase in synapse function, especially but not uniquely GABAergic.

### Gaucher disease organoids display alterations in excitatory and inhibitory neuronal function leading to network hyperexcitation

The results from lipid analyses, where hSOs displayed increased levels of lipids, and from single-cell RNA sequencing experiments, indicating a switch from GABAergic towards glutamatergic lineage development, prompted us to investigate functional deficits of inhibitory neurons in Gaucher disease. For this, we performed MEA recordings from our hCOs and hSOs using a 64-electrode MEA system and placing the organoids on the electrodes for acute recordings at DIV150 (Fig 5A). To examine potential GABAergic neuron deficits, we designed a paradigm to specifically identify deficiencies in this population by first recording basal activity, then adding PTX to the media to block inhibitory signalling and identify changes in excitation and finally adding NBQX and APV to block excitation (Fig 5A). We first verified that the paradigm was working as expected and that we could detect basal activity and increased excitation after PTX addition. We also confirmed the lack of activity following NBQX and APV addition, indicating NMDAR and AMPAR-dependent glutamatergic synaptic signalling is present (Fig 5B).

**Figure 5.**
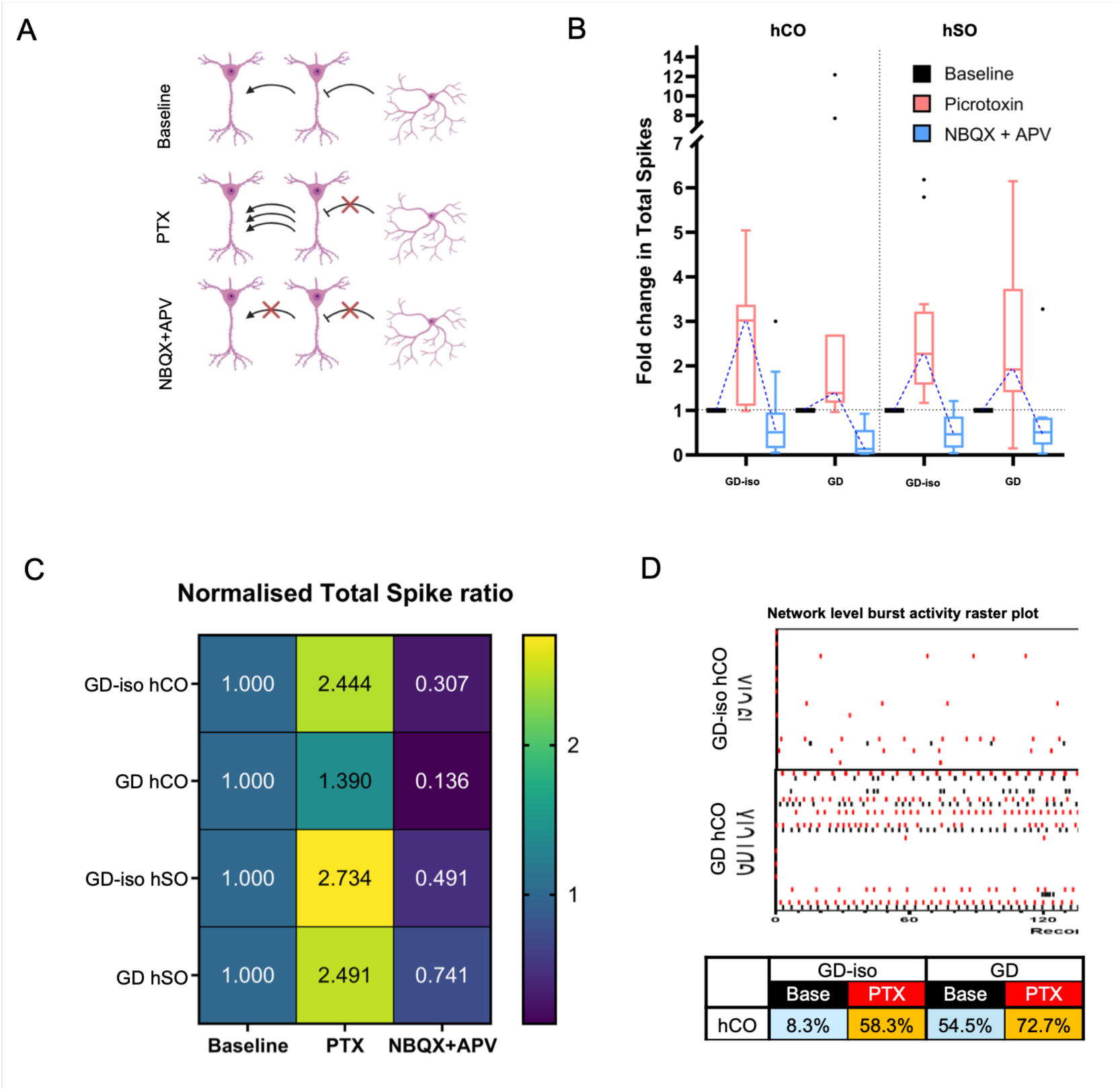
MEA recordings in brain organoids show a hyperexcitability phenotype of Gaucher disease hCOs. **A)** Schematic representation of organoid recording and experimental paradigm with an initial baseline recording, addition of picrotoxin (PTX) to continue recording, and addition of NBQX and APV to continue recording. **B)** Fold-change in total spike number of hCO and hSO at baseline (black), after PTX addition (red) and after NBQX and APV addition (blue), for GD-iso and GD organoids at DIV150. **C)** Normalised total spike ratio for each type of organoid (hCO and hSO) generated from the GD-iso and GD lines comparing PTX or NBQX + APV treatment to the baseline recordings (bottom graph). **D)** Raster plot showing burst activity detection from individual hCOs from GD-iso and GD lines at baseline (black) or after PTX addition (red) for a 5 min recording, with the percentage of active electrodes illustrated in the table for each condition.

After recording from 9-10 organoids from each line and fate, we observed, as expected, that when blocking GABA_A_R-dependent inhibitory signalling by adding PTX, we detected an increase in excitation for all our samples (Fig 5B). However, this increase was substantially different when comparing GD and GD-iso hCOs. While GD-iso had a 2.44-fold increase in spike ratio, GD showed a subtle 1.39-fold increase (Fig 5C). These results suggest that either inhibitory signalling is impaired in basal conditions and those organoids are less responsive to PTX, or the increase in glutamatergic vs GABAergic neurons triggers a hyperexcitation phenotype in Gaucher disease. The basal network burst activity of the GD hCOs was already much higher than that of the GD-iso control (Fig 5D), supporting the developmental switch as the main reason for the hyperexcitation of Gaucher disease organoids but without discarding alterations in GABAergic neuronal function. Conversely, when comparing HC to HC-KO (Fig S6A), the later had a 1.74-fold increase versus a 1.41-fold increase of HC.

For hSOs, changes in activity after PTX treatment were similar in both GD-iso and GD lines (Fig 5B,C) with a 2.73-fold and a 2.49-fold increase in spike ratio, respectively. Meanwhile, for the HC and HC-KO pair, the later had a much higher increase, with a 3.24-fold increase compared to the 2.03-fold increase of the HC (Fig S6A,B). As shown in the single-cell RNA sequencing experiments, hSOs are formed basically by GABAergic neurons, therefore, blocking inhibition had a major effect in facilitating activity within the network by disinhibition. In contrast to hCOs, when observing basal activity, we did not detect high activity levels in Gaucher compared to control organoids (Fig S6C), further suggesting that glutamatergic neurons are overexcitable and could be the driver of the hyperexcitation phenotype. Altogether, our results indicate that Gaucher disease cortical organoids have a higher basal activity and, therefore, are less responsive to network disinhibition, causing a hyperexcitation state of the neuronal network within organoids.

### Gaucher disease 2D cultures containing GABAergic and glutamatergic neurons identify cell-type specific contributions to network dysfunction

To help clarifying whether the developmental switch and different ratio of glutamatergic and GABAergic neurons is the sole cause of the hyperexcitation of Gaucher disease organoids, we performed additional high-density MEA recordings in 2D cultures. In these experiments, we focused on the use of the GD and GD-iso lines, being the more representative of Gaucher disease patients and a healthy control individual, with the advantage of being isogenic and therefore, share the exact same genetic background. Here, we leveraged forward programming strategies to independently generate pure populations of glutamatergic neurons, GABAergic neurons, and astrocytes. We then established GD and GD-iso cultures as well as chimeric cultures in which we mixed either healthy glutamatergic neurons and astrocytes with disease GABAergic neurons, or disease glutamatergic neurons and astrocytes with healthy GABAergic neurons (Fig 6A). With this approach, we were able to match the number of cells for each neuronal type and pinpoint their individual contribution to the network hyperexcitation independently of the developmental switch. In addition, this setting allowed us to perform weekly recordings of the cultures, from DIV14 to DIV54, to investigate network dysfunction over time. As expected, we saw a progressive increase in the median firing rate of neuronal activity over time for all conditions (Fig 6B), which started at around 0.5 Hz and ended between 0.76 and 1.55 Hz depending on the sample (Fig 6C). In line with organoid recordings, cultures with all cell types from the GD line presented a clear hyperexcitation of the network that started around DIV30 and increased with time in culture, with DIV50 cultures displaying a median firing rate of 1.55 Hz (Fig 6C). In contrast, cultures with all cell types from the GD-iso line had much lower excitation with the median firing rate at 0.76 Hz (Fig 6C). Interestingly, cultures with only disease GABAergic neurons performed very similarly to control cultures with a median firing rate at DIV50 of 0.86 Hz, while cultures with disease glutamatergic neurons but healthy GABAergic neurons displayed mild hyperexcitation with a median firing rate at DIV50 of 1.2 Hz (Fig 6C). For all conditions, both spike amplitude and median inter-spike interval did not differ significantly between conditions (Fig 6C). Upon treatment with PTX, the healthy glutamatergic and disease GABAergic condition showed a small tendency towards increased activity, suggesting potential hyperexcitability in GABAergic neurons that could lead to a higher suppression of glutamatergic signalling (Fig S6D). These results suggest that glutamatergic and to a smaller extent, GABAergic neurons display hyperexcitability, and dysregulation of the lineages may tip excitatory/inhibitory balance in Gaucher disease. However, as it was not possible to distinguish if firing comes from excitatory or inhibitory neurons, we complemented these analyses with calcium imaging recordings.

**Figure 6.**
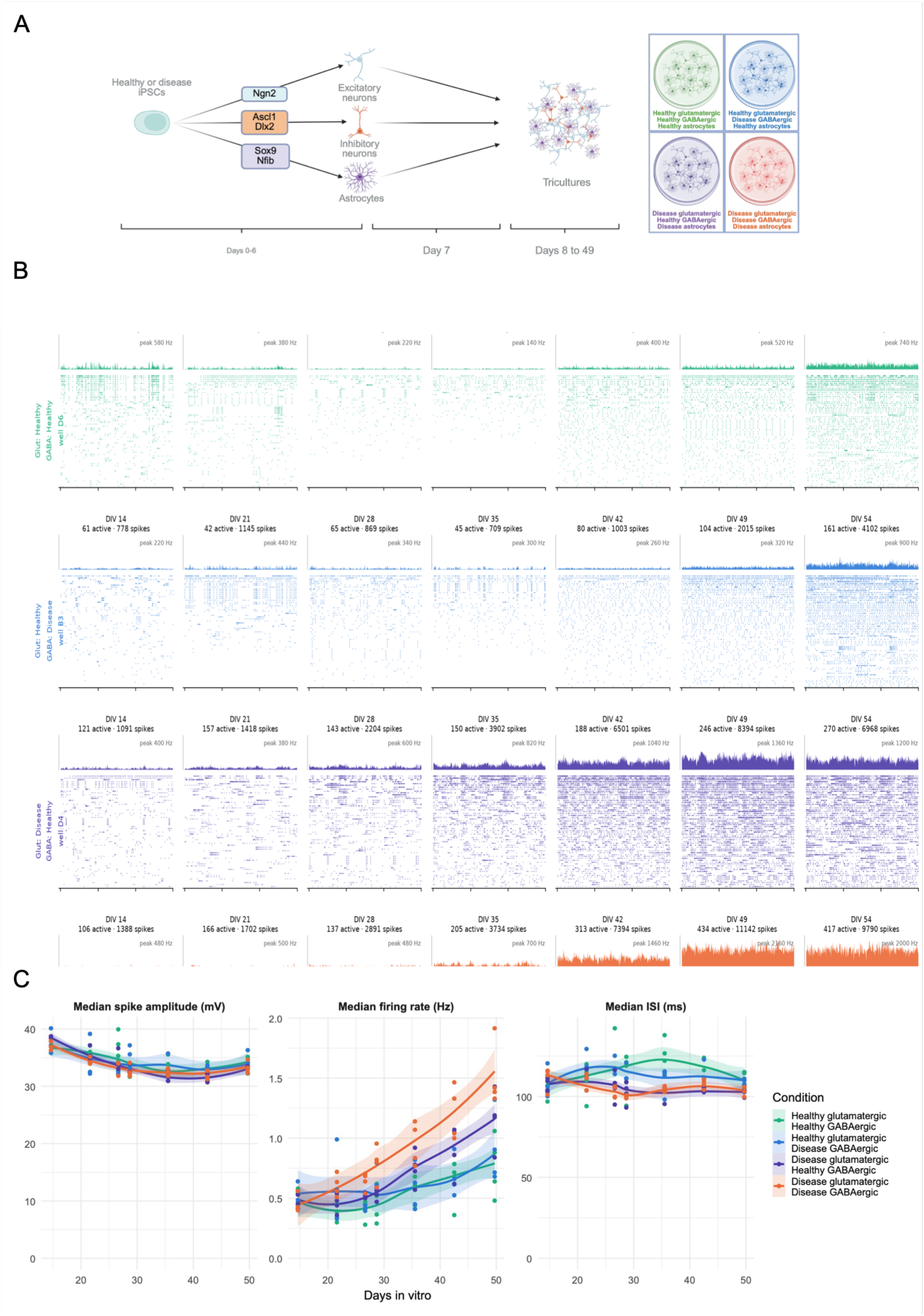
HD-MEA recordings in chimeric forward programmed tricultures show a hyperexcitability phenotype of Gaucher disease glutamatergic and GABAergic neurons. **A)** Schematic representation of the generation of tricultures consisting of excitatory neurons (Ngn2), inhibitory neurons (Ascl1 + Dlx2) and astrocytes (Nfib + Sox9) using transcription factor induction and establishment of chimeric cultures combining healthy/disease excitatory/inhibitory neurons. **B)** Raster plots showing spikes in one HD-MEA well of each culture condition over time, from DIV14 to DIV54. **C)** Graphs displaying the median spike amplitude (mV), median firing rate (Hz) and Median ISI (ms) of all coculture combinations over time.

For that purpose, we used a genetically encoded calcium indicator driven with a CAMKIIɑ promoter, providing expression specificity to glutamatergic neurons, recording first basal activity and later applying bicuculline (BIC) treatment to block GABA_A_R-dependent inhibition, a similar paradigm as our previous MEA recordings. Recordings were performed weekly across four weeks (DIV28, DIV35, DIV42 and DIV49), with a clear effect of BIC treatment on synchronising neuronal activity as shown by neuronal traces (Fig S7A) and raster plots (Fig 7A), and similar fraction of active cells across conditions (Fig S7B). From these experiments, we extracted and analysed the size of collective events by means of total number of cells participating in the events (Fig 7B), the number of peaks per cells (Fig 7C) and the area under the curve for each peak and cell (Fig 7D). Untreated cultures did not show major differences, except at DIV42 and DIV49 in the disease glutamatergic and healthy GABAergic conditions, where more cells became involved in collective events (63-70% compared to 48-57% in other conditions), suggesting a shift in the excitatory/inhibitory balance (Fig 7B). As expected, BIC treatment promoted an increase in the number of cells participating in collective events across conditions and timepoints (from 39-70% to 58-99% overall), however, increases were much more modest in conditions with disease glutamatergic neurons (always below 90.5%), especially if GABAergic neurons were also affected (below 75.8% for all timepoints, Fig 7B). Notably, the healthy glutamatergic and disease GABAergic condition showed a higher level of synchronisation after BIC treatment, with most cells (always above 97% from DIV35) being involved in collective events (Fig 7B). Notably, the fraction of cells contributing to populations events under baseline conditions was lowest at DIV28 and showed a trend of increasing over time, with increases occurring upon BIC treatment in all populations but less pronounced when containing disease glutamatergic neurons with healthy or disease GABAergic neurons (Fig S7C).

**Figure 7.**
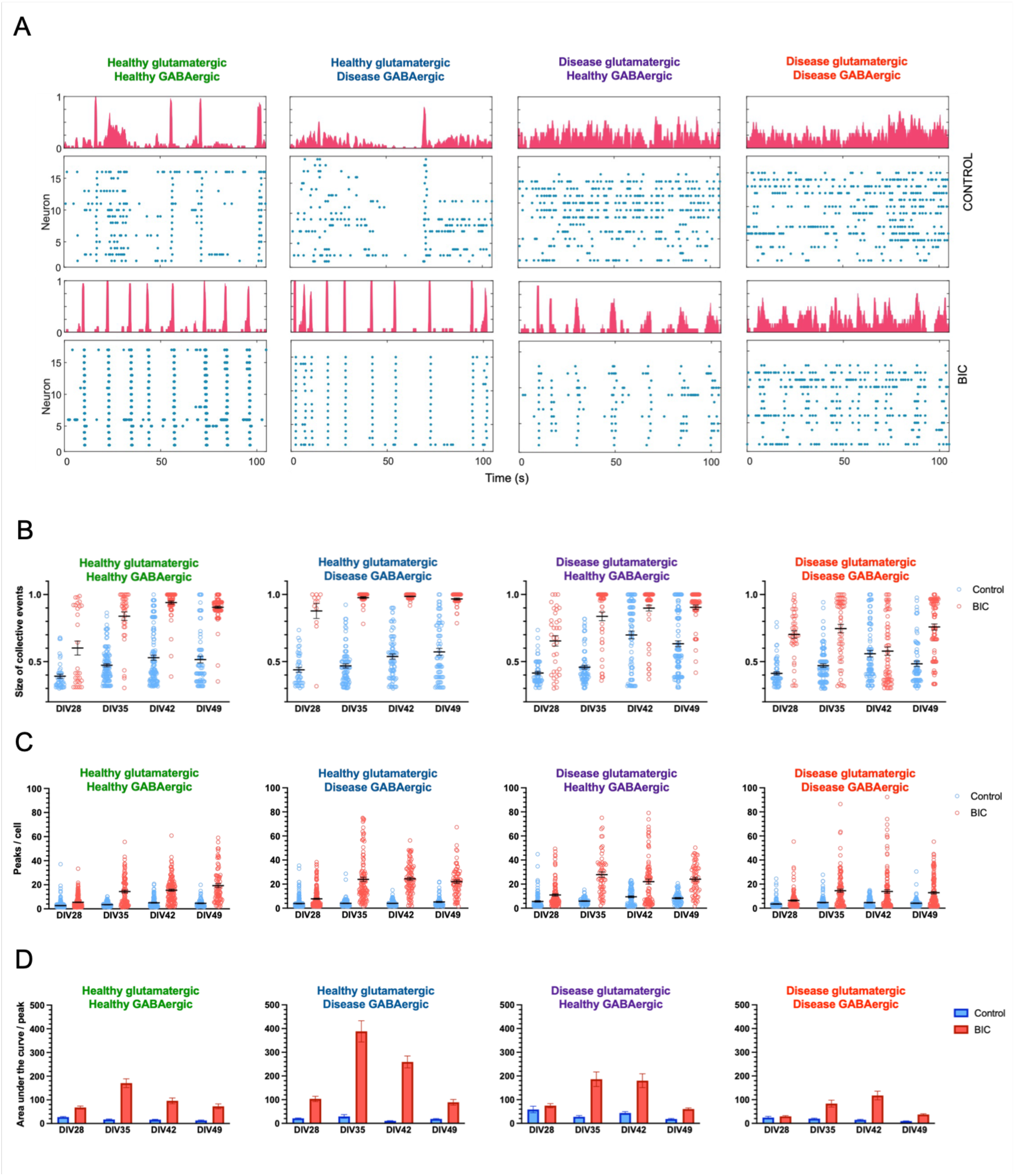
Calcium imaging of CAMKIIa-GCaMP8m in chimeric forward programmed tricultures show an excitation-inhibition imbalance. **A)** Raster plots showing spikes from a representation calcium imaging recording of each culture condition at DIV49 from the baseline (CONTROL) and immediately after bicuculline addition (BIC). **B)** Graphs displaying the size of collective events over time (from DIV28 to DIV49) before (Control) and after (BIC) bicuculline addition expressed as percentage of cells participating on collective activity events. **C)** Graphs displaying number of peaks per cell (from DIV28 to DIV49) before (Control) and after (BIC) bicuculline addition. **D)** Graphs showing the area under the curve of each peak over time (from DIV28 to DIV49) before (Control) and after (BIC) bicuculline addition.

The number of peaks per cell were also increased after bicuculline treatment, especially at DIV35, DIV42 and DIV49 with the chimeric conditions (healthy glutamatergic + disease GABAergic; disease glutamatergic + healthy GABAergic) showing higher increases and always above 22 peaks, while glutamatergic + GABAergic disease cultures showed the lowest increase, always below 14 peaks (Fig 7C). Upon examining the mean number of transients across experiments (Fig S7D), a significant increase was observed in baseline activity of disease glutamatergic cultured with healthy GABAergic neurons compared to healthy glutamatergic cultured with healthy GABAergic neurons at DIV28 (10.9 versus 7.7), at DIV35 (12.9 versus 10.2) and at DIV49 (11.7 versus 6.5) (Fig S7D). Differences were also significant comparing disease glutamatergic cultured with healthy GABAergic neurons to healthy glutamatergic cultured with disease GABAergic neurons at DIV35 (12.9 versus 9.9) and at DIV49 (11.7 versus 8.0) (Fig S7D). As the disease glutamatergic cultured with disease GABAergic neurons did not show the highest mean activity or a significant difference to the healthy glutamatergic cultured with healthy GABAergic neurons, this indicates a discrepancy between spiking frequency observed in network level recordings in MEA and examining an exclusively excitatory input, suggesting the remaining spiking increase could be caused by CAMKIIa-GABAergic neurons in response to hyperexcitable glutamatergic cells. Interestingly, the area under the curve of the peaks was clearly increased in the cultures with only disease GABAergic neurons, especially at DIV35 and DIV42 (388 and 259, respectively), showed an intermediate phenotype in cultures with only disease glutamatergic neurons (187 and 180, respectively) with lowest values in disease cultures (84 and 118, Fig 7D). This suggests that healthy glutamatergic neurons present homeostatic excitability correction to compensate for a stronger inhibition provided by disease GABAergic neurons, and a potential hyperexcitable state of the later, a mechanism that might fail in disease glutamatergic neurons. Collectively, calcium imaging experiments point towards GABAergic neurons being intrinsically hyperexcitable driving healthy glutamatergic neuron homeostasis to a more excitable state, which is revealed when removing inhibition.

Altogether, our results from MEA recordings and calcium imaging indicate that in Gaucher disease, glutamatergic neurons are clearly hyperexcitable. Meanwhile, GABAergic neurons whilst likely also hyperexcitable, seems to have a lower contribution to network alterations, with a limited capacity to buffer hyperexcitability, as shown by the fact that healthy GABAergic neurons cocultured with disease glutamatergic neurons display milder network hyperexcitability.

## DISCUSSION

Here, we established, to our knowledge, the first human forebrain iPSC-derived organoid model of Gaucher disease by generating both cortical and subpallial organoids, resembling the dorsal and ventral forebrain, respectively. We first generated a patient iPSC line (GD) and used it together with our previously established healthy carrier iPSC line (HC) to generate their corresponding isogenic lines using genome editing (GD-iso and HC-KO, respectively). Using this complete set of human iPSCs, we generated forebrain cortical and subpallial organoids (hCOs and hSOs) and demonstrated that our models recapitulated the lipid accumulation that occurs in patients. We decided to generate forebrain organoids because of the known neuropathological findings in patient brains indicating degeneration of cortical layers accompanied with gliosis^41^. Moreover, we showed a progressive accumulation of glycosphingolipids and a subsequent impairment in neuron development and maturation, leading to dysfunction of both glutamatergic and GABAergic neurons and a high overall network hyperexcitability.

The main hallmark of lysosomal storage disorders is the impairment of lysosomal function and accumulation of substrates, leading to an increase in number and size of lysosomes. However, previous reports using iPSC-derived neuronal models showed disparity of results, identifying either an increase^23, 24^ or a decrease^22, 26, 28^ in lysosomes, which can be due to the different markers used for the quantification (either lysosomal proteins or dyes). We used Lysotracker dye and identified clear increases in the HC-KO compared to the HC, but not in the GD compared to the GD-iso. In addition, we quantified GlcCer, the metabolite accumulating in Gaucher disease, and GlcSph, a secondary metabolite that serves as a biomarker. For GlcCer and similarly to Lysotracker, differences were only detected for the HC-KO organoids, while GlcSph was clearly elevated in both disease lines, although at a much higher level in the HC-KO. This is consistent with the differences in enzymatic activity, with the HC-KO line displaying around 60% of the activity compared to the GD line, and thus potentially being indicative of a perinatal lethal phenotype present in a rare few identified Type II Gaucher cases^42^.

Dysregulation of GlcCer metabolism leads to changes in complex glycosphingolipids for which GlcCer is the precursor, such as gangliosides, as shown in Gaucher disease patients^43^, in *GBA1*-related Parkinson’s disease patients^44^, and in cultured fibroblasts of type II Gaucher patients^45^. Lipid accumulation in our models appeared before there were clear changes in lysosomes, at least for the GD line (patient-derived), and it led to increased levels of GM_1_ membrane ganglioside, which were more evident in hSOs containing only inhibitory neuronal lineages, and suggesting higher susceptibility of these populations.

Gaucher disease can present in early-onset and severe neuronopathic forms^5, 6^, which hints towards the idea that prenatal brain development can be affected and trigger some of the later neurological manifestations. Importantly, gangliosides are essential for many developmental processes in the brain^38^, supporting the idea that prenatal brain development can be affected and lead to posterior neuropathological alterations. From the single-cell transcriptomics data of our brain models, we observed a striking early alteration of mitochondrial function, suggesting that either mitochondrial impairments play an important role in the early brain development pathology of Gaucher disease or an increase in differentiation and maturation of neurons with a consequent switch from glycolysis to oxidative phosphorylation. To support the first idea, mitochondrial impairment has been shown to be a common disease mechanisms in most LSDs, including Gaucher disease^46^. Moreover, a recent report identified a role of GCase in maintaining mitochondrial complex I function after being imported into mitochondria using iPSC-derived neuronal and midbrain organoid models^27^, supporting a role of GCase in mitochondria as an unrelated disease mechanism to glycosphingolipid accumulation. To support the second idea, we showed an increase in mitochondria TMRM staining in both glutamatergic and GABAergic neurons derived from GD lines compared to GD-iso, and our transcriptomics data for the late timepoint clearly showed a downregulation of progenitor identity and pathways related to proliferation, together with a striking upregulation of neuronal maturation and synaptic function pathways. In addition, mitochondrial alterations can be related to a potential interneuron vulnerability in GD, as fast-spiking interneurons exhibit higher oxygen consumption rates and are more sensitive to metabolic stress^47^.

Other cell types that rely on proper mitochondrial function are migratory cells. In subpallial organoids we identified a striking increase in the number of migrating interneurons, with a clear upregulation in pathways related to cholesterol metabolism. The dysregulation of cholesterol metabolism has been shown in Gaucher disease macrophages^48^ and it is known to play a role in *GBA1*-caused Parkinson’s disease^49^. Gangliosides and cholesterol are two of the main components in lipid rafts^50^, and changes in their abundance can alter the formation and number of these lipid rafts with the consequent changes in membrane receptors, including Trk receptors essential for cell survival^51^. Migrating neurons highly rely on signals from their environment, and these changes could promote aberrant or stalled migration with a subsequent accumulation of these cells. In addition, these cells present a striking downregulation of the Golgi apparatus, and Golgi alterations have been shown for other LSD and neurodegenerative disorders^52^, while proper Golgi function and polarity is essential for cellular migration^53^. Likewise, GM1 accumulation at the mitocohondrial-ER junctions has been shown to overload mitochondrial calcium and lead to heightened mitochondrial membrane permeabilisation^54^, and at the ER-plasma membrane junctions it induces synaptic spine formation^55^, supporting a hyperexcitabilty phenotype.

For our cortical organoids, the late stages of differentiation revealed an interesting switch from GABAergic to glutamatergic neurons in Gaucher disease organoids, even overcompensating the initial low number of glutamatergic neurons in patient organoids. Functional assays were performed to investigate the effect of this switch on network functionality, showing a clear hyperexcitation of the network in Gaucher disease. Many neuronopathic Gaucher disease patients present seizures^56^, which are most likely caused by a hyperexcitability of the neuronal networks due to excitation/inhibition imbalances^57^. Our results support this hypothesis; however, they do not allow to pinpoint the exact contribution of excitatory versus inhibitory neurons to this phenotype. Previous studies have shown a higher resting membrane potential of iPSC-derived neurons from Gaucher patients^21^, suggesting a higher excitability level of glutamatergic neurons. To fully explore glutamatergic and GABAergic dysfunction in our models, we independently generated glutamatergic and GABAergic neurons from the GD and GD-iso lines and established chimeric cultures with all possible combinations of healthy and diseased neurons. HD-MEA and calcium imaging recordings demonstrated that glutamatergic neurons are hyperexcitable, while hyperexcitable GABAergic neurons still fail to compensate for this hyperexcitability, but can further supress healthy glutamatergic neurons. This is an interesting finding, since our transcriptomic data points towards an increase in GABAergic synapse function, which could be explained by an attempt of the organoid to compensate for the hyperexcitability and loss of GABAergic neurons, which in addition are less efficient to control hyperexcitation of glutamatergic neurons. It remains to be clarified whether the increase in genes related to GABAergic synapses is matched by an actual increase in synapse numbers or function, which could be targeted towards other GABAergic neurons reducing their efficiency to decrease excitation of glutamatergic neurons.

In conclusion, we have generated a novel brain organoid model of Gaucher disease and identified disease hallmarks, lipid dysregulations and a subsequent alteration in neuronal lineage development and maturation leading to neuronal network dysfunction, which can explain some of the neuropathological findings identified in Gaucher disease patients. We anticipate these models will be useful to continue investigating disease mechanisms and identify potential targets for therapeutic development to treat this devastating childhood disorder.

## MATERIALS AND METHODS

### Fibroblast cultures

Human skin fibroblasts (GM08398, GM01260) were obtained from the Coriell Institute for Medical Research by material transfer agreement. Fibroblasts were maintained in Dulbecco’s modified Eagle’s media (DMEM; Thermo Fisher Scientific) supplemented with 10% Fetal Bovine Serum (FBS; Thermo Fisher Scientific) and 1% GlutaMAX (Thermo Fisher Scientific), at 37°C in humidified air with 5% CO_2_, and passaged upon reaching ∼80% confluency using 0.25% Trypsin (Thermo Fisher Scientific).

### Generation and maintenance of iPSC lines

Human iPSCs were generated by the Cell & Gene Therapy Core Facility at Lund University through fibroblast mRNA-based reprogramming in accordance with the StemRNA™ 3rd Gen Reprogramming Kit for Reprogramming Adult and Neonatal Human Fibroblasts (Stemgent®; ReproCell) protocol. Clonal iPSC lines were established by manually selecting human ESC-like colonies and characterised in accordance with the recommended Standards for Human Stem Cell Use in Research set out by the International Society for Stem Cell Research^35^.

Human iPSC lines were maintained in mTeSR1 (StemCell Technologies) and passaged by dissociation with StemPro Accutase Cell Dissociation Reagent (Thermo Fisher Scientific) upon reaching 80% confluency. After collection, cells were centrifuged at 300 x g, and replated at a density of 1.5 x 10^4^–2.5 x 10^4^ cells / cm^2^ in mTeSR1 supplemented with 10 µM Rho kinase inhibitor Y-27632 (RI; Tocris Bioscience) in 6-well plates coated with hESC-qualified Matrigel (Corning). Cultures were then left overnight in a 37°C, 95% O_2_ and 5% CO_2_ incubator, followed by a full daily media exchange with fresh mTeSR1. Routine mycoplasma testing of all lines was performed using MycoAlert Mycoplasma Detection kit (Lonza).

### GBA1 mutation analysis

*GBA1* genotype was confirmed by performing genomic DNA extractions (DNeasy Blood & Tissue kit, QIAGEN) of fibroblasts and iPSC lines, subsequent PCR of all *GBA1 exons* (Table S2; Primers, Table S3; PCR settings) using a Master Cycler Nexus Thermocycler (Eppendorf) with Phusion DNA polymerase (Thermo Fisher Scientific) in HF buffer following the manufacturer’s instructions. Sanger sequencing was performed using the service of EuroFins Genomics and sequences were aligned in SnapGene (V5.0) to the Ensembl genome browser reference genome sequence for human *GBA1* gene (ENSG00000177628).

### G-band and molecular karyotyping

G-Band karyotypic sample preparation was performed as previously reported^58^, and karyotypic analysis was performed at Sant Joan de Déu Barcelona Children’s Hospital (Spain). Molecular Karyotyping was assessed using copy number via SNP-array from genomic DNA with IlluminaBeadArray Technology at LIFE & BRAIN GmbH via Infinium Global Screening Array-24 BeadChip (Illumina). Genotype/CMV analysis via GenomeStudio V2.0.5 and CNV-Partition 3.2.0 was performed as a service by the Cell & Gene Therapy Core Facility at Lund University. Minimum probe count was set to 30, reportable cut-off was set to 400 kb for gains/losses and 5 Mb for loss of heterozygosity regions with a confidence cut off at 35.

### Immunocytochemistry

iPSC colonies were cultured in 12-well chamber slides (Ibidi GmbH) and then fixed in 4% paraformaldehyde in PBS (PFA; Electron Microscopy Sciences) for 15 min at room temperature and washed thrice with phosphate-buffered saline (PBS; Thermo Fisher Scientific). Fixed cells were blocked and permeabilized with PBS containing 0.025-0.25% (for extracellular or intracellular epitopes, respectively) Triton X-100 (Electron Microscopy Sciences) and 5% normal donkey serum (Sigma-Aldrich; blocking solution) for 1 h at room temperature. Primary antibodies were incubated overnight (ON) in blocking solution at 4°C. The following day, cells were washed twice for 5 min with PBS and once for 5 min with blocking solution. Cells were incubated with blocking solution containing secondary antibodies and Hoescht 33342 (2 µg/µl; Thermo Fisher Scientific) for 2 h at room temperature, then washed three times with PBS for 5 min before being stored at 4°C protected from light. All antibodies and working dilutions can be found in Table S4. Images were acquired with a 20x objective on a Leica SP8 inverse confocal using LAX software and prepared in Fiji.

### Reverse transcription quantitative PCR (RT-qPCR)

Determination of pluripotency gene expression in iPSC clones was achieved via RT-qPCR. RNA extraction was carried out directly from plated ∼80% confluent iPSCs from three independent passages using the RNeasy Mini Kit (QIAGEN) according to the manufacturer’s instructions and with treatment of RNA with DNase I (QIAGEN) to prevent DNA contamination. Eluted RNA concentration was quantified using a Nanodrop One spectrophotometer (Thermo Fisher Scientific), and subsequently 1 µg of RNA was used in a reverse transcription reaction using the qScript cDNA Synthesis Kit (QuantaBio) following manufacturer’s instructions alongside negative controls without transcriptase. RT-PCRs were carried out in a final volume of 15 µl with 3 technical replicates and 1 negative control per sample, using TaqMan™ Universal PCR Master Mix (Thermo Fisher Scientific) and TaqMan™ Real-Time PCR assays (Thermo Fisher Scientific; listed in Table S5) following manufacturer’s instructions for 40 cycles on a QuantStudio 7 system using Design and Analysis 2.6.0 software (Bio-Rad).

### Trilineage differentiation

Human iPSCs were plated onto Matrigel-coated 24-well µ-plates (Ibidi GmbH) with mTeSR1 media supplemented with 10 μM of RI at a density of 1 x 10^5^ cells per well for endodermal and ectodermal fates, and 5 x 10^4^ for mesodermal fate. Differentiation towards the three germ layers was performed using STEMdiff™ Trilineage Differentiation Kit (StemCell Technologies) for seven days, before being fixed with 4% PFA for 15 min and immunostained for markers of corresponding germ layer. All antibodies and working dilutions can be found in Table S4. Images were acquired with a 20x objective on a Leica SP8 inverse confocal using LAX software and prepared in Fiji.

### Generation of genome editing constructs

For standard CRISPR/Cas9, two sgRNAs were designed (Table S6) using the Benchling CRISPR Guide RNA design tool (www.benchling.com) the sequence. For selectivity for the *GBA1* gene loci over the *GBA1LP* pseudogene, sgRNA-4 was design to target a 55-bp non-homologous region in exon 9 of the *GBA1* gene. Designed oligonucleotides were synthesised (Eurofins Genomics), and 1 µl single stranded forward and reverse oligonucleotides (100 mM) which are partially complementary were phosphorylated and annealed in a solution of 1 µl 10x T4 DNA ligase reaction buffer, 0.5 µl T4 Polynucleotide Kinase and 6.5 µl ddH_2_O. The reaction was performed in a thermocycler with 37°C for 30 min, then 95°C for 5 min, before ramping down to 25°C at 5°C / min. The oligonucleotide duplex was then diluted 1:250 in ddH_2_O for later steps. A digestion-ligation reaction was performed with 100 ng of pSpCas9n(BB)-2A-Puro (pX459) V2.0 (a gift from Feng Zhang, Addgene #62988) or pSpCas9n(BB)-2A-Puro (pX462) V2.0, (Addgene plasmid #62987), diluted into a solution containing 2 µl of the oligo duplex, 2 µl 10x FastDigest Buffer, 1 µl DTT (10 mM), 1 µl ATP (10 mM), 1 µl FastDigest Bpil, 0.5 µl T7 DNA ligase, adjusted with ddH_2_O to a final reaction volume of 20 µl. The reaction was performed in a thermocycler with 6 cycles of 37°C for 5 min followed by 23°C for 5 min. An 11 µl volume of the reaction was then treated with 1 µl of Plasmid-safe DNase with 1.5 µl ATP (10 mM) and 1.5 µl 10x Plasmid-Safe buffer and incubated at 37°C for 30 min. A single-strand donor oligonucleotides (ssODNs) for the *GBA1* P415R correction was designed to contain 30 nt homology arms on each side of the sgRNA target sequence, a silent mutation in the PAM sequence (red nt in Table S6), and phosphothiorate bonds at both ends of the donor strand (* in Table S6). The ssODN was synthesized by Integrated DNA Technologies.

For Prime Editing genome editing, the *GBA1* exon 10 containing the L444P mutation was targeted. The pegRNAs and ngRNAs design candidates for PE3b Prime Editing were identified using PrimeDesign^59^. The pegRNAs were chosen depending on the length of the primer binding site and the retrotranscriptase template, prioritizing those in which the targeted PAM site was disrupted by a silent mutation. For the PE3 ngRNA, those that incorporate the edited nucleotides (PAM silencing and mutation correction) were chosen as optimal, with a focus on those that had both edits (PE3b) to enhance the efficiency of nicking the non-edited strand (PE3b seed ngRNAs). All necessary sequences can be found in Table S7. All oligonucleotides were ordered from EuroFins Genomics.

To create a pegRNA with our desired insert, 2 µg of the pU6-pegRNA-GG-Vector acceptor plasmid (Addgene: plasmid 132777) was first digested using BsaI restriction enzyme for 4 to 16 hr at 37°C before being isolated from its insert via electrophoresis of a 1% agarose gel, followed by gel excision and DNA extraction using the Gel Extraction Kit (QIAGEN) following manufacturer’s instructions. The complementary top and bottom oligonucleotides pegRNA spacer, the pegRNA extension, and the scaffold were individually prepared in annealing buffer, composed of Tris-Cl (10 mM) and NaCl (50 mM). Samples were incubated at 95°C for 3 min then cooled gradually (0,1°C / s) to 22°C, before being diluted to a final concentration of 1 µM for each oligonucleotide. The digested plasmid along with the annealed oligonucleotides (spacer, extension and scaffold) were assembled with Ligase T4 enzymes in a buffer containing BsaI-HF v2 to prevent ligation of the empty vector. The mix was incubated 10 min at room temperature, 15 min at 37°C, 15 min at 80°C before being held at 12°C. Finally, the plasmid was transformed in competent cells and antibiotic-resistant clones were picked and analysed by sequencing to determine if the insert was correctly inserted.

To create ngRNAs for PE3b, oligonucleotides for the top and bottom of the sgRNA were annealed using T4 Polynucleotide Kinase at 37°C for 30 min, 95°C for 5 min and then cool gradually (5°C/min) to 25°C. The plasmid BPK1520 (Addgene: Plasmid #65777) and annealed sgRNA oligos were ligated together using T7 DNA ligase and fast digest enzyme Esp3I to cut the plasmid via incubating the mix at 5 min at 37°C and 5 min at 23°C for 6 cycles. To remove any linear DNA, the plasmid was incubated for 30 min at 37°C with Plasmid-Safe DNase and ATP. As a final step, all plasmids were transformed in competent cells and the positive clones were picked and analysed by sequencing.

To determine if the correct sequences were integrated, plasmids were isolated using the Plasmid Mini Kit (QIAGEN) following manufacturer’s instructions. The purified plasmids were sequenced using the hU6 primer by EuroFins Genomics. Sequences were aligned against the predicted sequence generated in SnapGene Software.

### Genome editing of patient iPSC lines via CRISPR-Cas9 and Prime Editing

For high-yield plasmid preparation to transfect cells, plasmids were purified using the EndoFree Maxi Kit (QIAGAEN) following manufacturer’s instructions. DNA quantifications were done with a NanoDrop 1000 Spectrophotometer and NanoDrop 1000 3.8.1 software.

To correct the P415R mutation from the GD disease patient iPSC line, CRISPR-Cas9, sgRNA and ssODN constructs were transduced into cells as follows to promote homology directed repair (HDR). On day -1, human iPSCs at ∼80% confluency were dissociated with Accutase, and 4,5 x 10^5^ cells were replated on Matrigel-coated 6-well plates using mTeSR1 medium supplemented with 10 µM RI. One day later (Day 0), medium was replaced with fresh mTeSR1 medium supplemented with 10 µM RI. Two separate tubes with 100 µl OptiMEM and 6 µl LipoStem, and 100 µl OptiMEM, 1.5 µg CRISPR-Cas9 and 0.5 µg ssODN, respectively, were prepared, mixed and incubated for 10 min at room temperature before being added to the cells. The two following days (Day 1 and 2), medium was replaced with fresh mTeSR1 medium supplemented with 10 µM RI and puromycin (1.25 ug / ml). On days 3 and 4, medium was replaced with fresh mTeSR1 medium supplemented with 10 µM RI. From day 5 medium was replaced with fresh mTeSR1 medium. When cells reached ∼50% confluency, they were dissociated with Accutase and 100, 200 and 1 x 10^4^ cells were plated per well on laminin-521-coated 6-well plates using mTeSR1 supplemented with 10 µM RI. For the following 3 days, medium was replaced with fresh mTeSR1 supplemented with 10 µM RI, with gradual decrease of RI concentration as individual colonies increase in size. Once individual colonies reached a diameter of 1-2 mm, they were picked and plated in individual wells of Matrigel-coated 24-well plates with mTeSR1 supplemented with 10 µM RI and PenStrep (100 µg/mL). Medium was changed daily for fresh mTeSR1 until cells reached ∼80% confluency. Cells were then dissociated using Accutase and split 1:4 before being plated on Matrigel-coated 24-well plates with 10 µM RI-supplemented mTeSR1. The remaining cells were pelleted for DNA extraction and subsequent genome editing assessment. To generate a *GBA1* knock-out line HC iPSCs the same sgRNA and protocols were utilised in the absence of ssODNs, in order to promote non-homologous end joining. The same approach was used to generate the KO line but using pX462, two sgRNAs and without the SSODN.

For nucleofection, human iPSCs of 60-70% confluence at passage number below 20 were used. Colonies were first rinsed with Dulbecco’s PBS (DPBS) without Ca and Mg, and then passaged by incubation in 1 volume (500 µl per well of a 6-well plate) of Accutase for 6 min at 37°C, generating a single-cell suspension. Accutase was neutralized with 4 volumes of DMEM. 8 x 10^5^ cells were counted, centrifuged at 300 x g for 5 min and resuspended in 90 µl P3 Lonza solution. For the PE3b version, 0.5 µg / µl of the plasmids at a ratio of 3.5:1:0.5 total DNA (PE-MAX, pegRNA and sgRNA constructs, respectively) were added to make up 5 ug in 10 µl to reach a total volume of 100 µl. The cell suspension was nucleofected in a nucleocuvette using the programme CB150 (Lonza 4D Nucleofector), and subsquently cells were plated in three wells of a 6-well plate coated with ES-Matrigel with mTeSR+ media and Clone R2 supplement (Stem Cell Technologies). The next 3 days, cells were fed with mTeSR^+^ and Blasticidin (10 µg / ml) to select only the transfected cells. On day 4, cells were passaged and serially diluted to reach 1000 cells / ml and plated at 40 cells per well of a 6-well plate with mTeSR+ media and Clone R2 supplement. When colonies reached a size of 1-2 mm, they were picked and transferred to 48-well plates for clonal expansion.

DNA was extracted from the frozen pellets using the Blood and Tissue Kit (QIAGEN), following manufacturer’s instructions. Quantifications were done with a NanoDrop 1000 Spectrophotometer and NanoDrop 1000 3.8.1 software. To confirm the correction of the mutation the PCR products were digested with MboI (for P415R) or with XmaI (for L444P) restriction enzyme following the manufacturer’s instructions (Thermo Fisher Scientific). Digested DNAs were run on an agarose gel containing 1x GelRed Nucleic Acid Stain (Biotium), in TAE 1x (3% agarose low melting temperature for the Exon 9-11 PCR and 1.5% gel agarose for Exon 8-10 PCR). To determine individual allelic genotype, Exon 8-10 and Exon 9-11 PCRs were performed, the resulting cDNA was utilised in Blunt End TOPO-cloning following the manufacturer’s instructions (Thermofisher Scientific) and using 4 µl of purified DNA. Transformation was performed in 50 µl competent OneShot TOP10 chemically competent E. Coli. To determine the allelic distribution 20 colonies were picked, expanded, purified using the Plasmid Mini Kit (QIAGEN) following manufacturer’s instructions, and sequenced with the hU6 primer by EuroFins Genomics. Sequences were aligned against the Ensembl human *GBA1* (ENSG00000177628) sequence in SnapGene Software.

### Off-target analysis

The top-5 most likely regions for off-target effects for the P415R sgRNA were predicted using Benchling software (Table S8), according to the aggregation scoring approach^60^ and analysed through PCR amplification of the region of interest using primers found in Table S9. Reactions were run using 2 min of initial denaturation, followed by 30 cycles of 98°C for 10 s, 66°C for 30 s, and 72°C for 30 s, prior to a final 72° step for 10 min. PCR products where then sequenced by EuroFins Genomics. Sequences were then compared against the corresponding sequence for the parental line.

### Enzymatic activity assay in induced neurons

Generation of induced neurons (iNs) was performed following a previously published protocol^61^. Briefly, iPSCs at ∼80% confluency were dissociated with Accutase (Day -2) and 3.5 x 10^5^ cells were plated on Matrigel Growth Factor Reduced Basement Membrane Matrix (Corning) coated 6-well plates with mTeSR1 supplemented with 10 µM RI. The following day (day -1), media was replaced with fresh mTeSR1 media, and 1 µl of rtTA and Ngn2 lentivirus were added to each well. Lentiviral vectors for FUW-M2-rtTA (reverse tetracycline-controlled transactivator, a gift from Rudolf Jaenisch (Addgene, #20342) and tetO.NGN2.puro a gift from Marius Wernig (Addgene, #52047) were prepared as previously described^62^. On day 0, media was replaced with fresh mTeSR1 media containing 2.5 µg / ml of doxycycline (Dox; Sigma-Aldrich), which was kept in the media throughout experiments. From day 1, BrainPhys media (StemCell Technologies) supplemented with 0.5% N2 and 1% B27 supplement was used, and 72 hr of 1.25 µg / ml puromycin selection was performed. From day 5, daily media exchanges were performed using BrainPhys supplemented with 0.5% N2, 1% B27, 10 ng / ml NT3 (Peprotech) and 10 ng / ml BDNF (Peprotech).

At day 7 iNs were rinsed with DPBS without Ca and Mg, dissociated with Accutase, pelleted at 300 x g and frozen at -80°C for enzymatic assay using a previously described method^63^ with minor modifications. Briefly, pellets were resuspended in McIlvain extraction buffer (0.2 M of Na_2_HPO_4_, 0.1 M of citric acid, 1% sodium taurate, 0.2% Triton X-100). Cells were then disrupted by sonication for 2 min at 10 s intervals of 100W (70 Amp) on ice. The lysates were centrifuged at 20000 x G for 20 min at 4°C to pellet insoluble proteins and the supernatant was transferred to a fresh tube on ice. A BCA assay (Thermo Fisher Scientific) was performed to normalize protein levels to 1 mg / ml. 20 µl of the normalised samples were added to 180 µl GCase buffer (62.86% 0.2 M Na2HPO4, 36.64% 0.1 M citric acid, 15 mM 4-methylumbelliferylglucopyranoside, 0.25% sodium taurocholate, 0.15% Triton X-100, 0.1% bovine serum albumin) to reach a final volume of 200 µl, and the mix was incubated for 1 hr at 37°C. After incubation, 800 µl of stop solution (500 ml: dH_2_O, 3.755 g glycine, 1.545 g NaOH, adjusted to pH 10.4) was added to end the reaction, and fluorescence of each sample was measured (ex. 365nm; em. 460nm) on a BioTek Cytation5 Cell Imaging Multimode reader (Agilent). One GCase activity unit (U) releases 1 nmol of 4-methylumbelliferone (4MU) per hr, and thus enzymatic activity was calculated using the following formula:

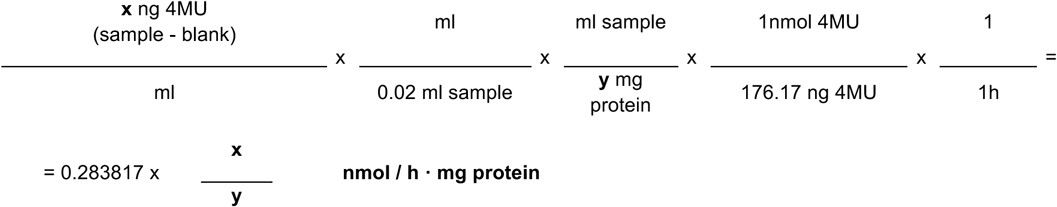

### Organoid culture

Human cortical organoids (hCOs) and subpallial organoids (hSOs) were prepared based on an adaptation of previously published protocols^37, 64^. Commercially available STEMdiff Dorsal and Ventral Forebrain Organoid Differentiation kits (StemCell Technologies) were used as recommended by manufacturer with minor modifications. iPSCs were grown to 60-70% confluency then dissociated, centrifuged at 300 x g for 5 min and resuspended at concentration of 3 x 10^5^ cells / ml in Seeding medium supplemented with 10 µM Y-27632 and plated into a Aggrewell 800 (StemCell Technologies) already containing 1 ml of Seeding medium. The Aggrewell was then centrifuged at 100 x g for 3 min to capture the iPSCs evenly into the microwells and incubated overnight to form initial organoid bodies. On days 1 – 5, there was a daily exchange of 1.5 ml fresh STEMdiff Neural Organoid Basal medium 1. On day 6, Aggrewell content was resuspended with a wide-bore 1 ml pipette tip, collected to a 37 µm reversible filter before being into a conical tube using freshly prepared Forebrain Organoid Expansion media (STEMdiff Neural Organoid Basal medium 2, Supplement A, Supplement B ± Supplement D; Supplement D for hSO only), and 40-50 organoids were plated per well into ultra-low attachment 6 well plates with 2 ml of medium and incubated for 48 hrs until day 8. Full medium changes with Forebrain Organoid Expansion medium were performed every 48 hrs from days 8 to 25. On day 25, organoids were redistributed to 15-20 organoids per well in ultra-low attachment 6 well plates, and medium was fully exchanged to Forebrain organoid differentiation media (STEMdiff Neural Organoid Basal medium 2, Supplement A, Supplement C), with full medium exchanges occurring every 2 days until day 43. On day 43, organoids were redistributed to 8-10 organoids per well in ultra-low attachment 6 well plates, and medium was fully exchanged to 3 ml Maintenance media (Neurobasal A, 2% B27-Vitamin A supplement, 1% Glutamax, 1% Penicillin/Streptomycin) and exchanged every 2 days until day 49. From day 49 onwards, organoids were transferred to an orbital shaker rotating at 80 rpm to improve tissue expansion and nutrient diffusion and 3 ml Maintenance media was exchanged every 2-3 days until organoids were processed.

### FACS stainings of organoids

Briefly, a single-cell dissociation was generated from organoids at the Day 75 and Day 150 timepoints as described below in “Single-cell RNA sequencing”. Following the final centrifugation pellets were resuspended in 900 µl chilled PBS and filtered through a 70 µm strainer before cell counting and normalisation of cell density between control and disease conditions. A 250 µl volume of suspension were transferred into a DNA LoBind Eppendorf per staining condition. A 250 µl volume of the 2x staining solution (1:5000 LysoTracker Deep Red, 1:100 LipidTOX Deep Red or 1:500 Lipidraft 555 CT-B conjugate) was then added to individual tubes for experimentation. Lipidraft 555 CT-B conjugate sample were incubated 10 min at 4°C before centrifugation at 200 g for 3 min, followed by resuspension in chilled PBS and a further 200 g for 3 min prior. The Lipidraft 555 samples were then resuspended in a 500 µl volume of 1:200 Lipidraft 555 CT-B antibody staining sample, and incubated at 4°C for another 15 min. Lysotracker DR and LipidTOX samples were incubated at room temperature for 30 min. Once staining was completed, all samples were centrifuged at 200g for 3 min then resuspended in 200 µl chilled PBS and treated with DAPI to a final concentration of 0.1 µg/ml to assess viability.

### Thin layer chromatography (TLC) of gangliosides

Lyophilized cells were dissolved in MeOH/CHCl_3_ (1 ml, 1:1 v/v), sonicated / homogenized (VWR Ultrasonic cleaner bath) for 30 min, and cooled to room temperature after sonication. Then Ultrapure H_2_O (0.5 ml) was added to the sample and sonicated for another 10 min. Gangliosides were separated from tissue and proteins by centrifugation at 2000 rpm (VWR MiniStar Silverline) for 20 min. The supernatant (upper phase, aquatic phase) was isolated, H_2_O (0.25 ml) added to the organic phase, and phase separation was induced by centrifugation for 20 min. The upper phase was isolated and passed through a Waters LC-18 column. Gangliosides were first eluted using methanol (6 ml) and then MeOH/CHCl_3_ (6 ml, 1:1 v/v). The fractions containing gangliosides were collected, transferred to an LC-MS vial and evaporated *in vacuo.* Total ganglioside fraction weight was obtained when the sample was fully desiccated. The resulting residue was then dissolved in MeOH/CHCl_3_ (1:1 v/v), analysed by TLC on 20 x 10 cm Silica F_254_ 15 µm glass plates using MeOH/CHCl_3_/0.2% CaCl_2_ (aq) (45:55:10) as mobile phase, developed using orcinol stain, and identified against authentic porcine brain ganglioside standards (Avanti Polar Lipids) by comparing Rf values of the sample gangliosides with the internal standard. The plates were photographed with a 48MP Sony IMX789 sensor (smartphone camera, OnePlus 10 Pro). Band intensities were quantified using ImageJ software.

To normalise TLC by total protein content, the combined tissue and protein solid phases were dissolved in 0.1M NaOH (500 µl final volume). The samples were then sub-aliquoted and diluted 1:4 with 0.1M NaOH before a Lowry assay was performed (Thermo Fisher Scientific) following manufacturer’s instructions.

### LC-MS/MS quantification of glucosylsphingosine (lyso-Gb1)

Day 75 organoids were collected, and supernatant was removed prior to flash-freezing on dry ice. Samples were weighed and approximately 100 µl ice-cold PBS was added per 10 mg tissue on ice. Samplers were then sonicated to generate a homogenous lysate, and 10% volume was removed to perform protein quantification via a BCA assay (Thermo Fisher Scientific). To the remaining sample, 3 volumes of ice-cold methanol were added and gently mixed to make a 1:3 PBS to methanol ratio. This mix was vortexed to homogenise, and centrifuged at 1500 x g for 5 minutes at 4°C before the supernatant was transferred to a fresh reaction tube, and this supernatant sample was then frozen until use. After thawing on ice, to 100 µl thawed sample, 20 µl of Internal Standard-Mix were added together with 300 µl of acetonitrile. The mixtures were vortexed for 3 min and incubated at –20°C for 20 min. Then, suspensions were centrifuged with a Centrifuge 5430 (Eppendorf) at 1300 rpm for 10 min. The whole supernatant was transferred into a new reaction tube and the remaining pellet was discarded. The supernatant was carefully evaporated under nitrogen gas using a Techne™ Sample Concentrator (Thermo Fisher Scientific) set to 40°C for 45 to 60 min. All samples were reconstituted using 100 µl of water/acetonitrile/formic acid 65/35/0.1. After reconstitution, samples were centrifuged at 1300 rpm for 10 min and 90 µl of the solution were transferred into a new LC-vial. 15 µl were injected on the column and analysed by LC-MS/MS.

### Single-cell RNA sequencing (scRNAseq)

Day 75 ± 3 (7-10 organoids) and day 150 ± (2-3 organoids) organoids were collected across 3 independent differentiations for initial dissociation. Dissociation was performed as previously reported^65^. In brief, Enzyme stock solution (ddH2O, 1x EBSS, 30% D(+)-Glucose, 1M NaHCO3, 50 mM EDTA) was equilibrated in a 5% CO2, 95% O2 atmosphere at least 12 h in advance. Depending on organoid age, 30 - 50 U / mL Papain was added to the Enzyme stock solution, followed by 4 mg of L-cysteine per 20 ml and 12500 units/ml DNase I, and then the solution was incubated at 37°C for 15 min before filtering. Organoids were transferred to a sterile Petri dish without liquid and a No. 10 blade (Swann Morton) was used to chop the organoid into 1 mm^2^ pieces over 30 s. After chopping, 10 ml of filtered Enzyme working solution was added to each petri dish, and the dishes were transferred to an orbital shaker in the cell culture incubator for 75 – 90 min. The solution containing the tissue pieces was then transferred to a LoBind DNA tube (Eppendorf) containing 4 ml Low Ovo solution before gentle centrifugation at 100 x g for 2 min. The supernatant was discarded, and the pellet was gently triturated 10-20 times with 1 ml Low Ovo solution until cloudy. The solution was allowed to settle for 30 s, then the supernatant was transferred to a new tube, and another 1 ml of Low Ovo solution was used to triturate the remaining tissue pieces. Next, 1 ml High Ovo solution was pipetted underneath the dissociated solutions and gently centrifuged at 100 x g for 5 min to separate live cells from debris. Cells were then resuspended in 0.04% human serum albumin in PBS and centrifuged at 350 x g for 5 min at 4°C to rinse the pellet.

From this point an adapted version of the Chromium Fixed RNA Profilling Sample Preparation kit (10X Genomics) was performed. Supernatant was removed and resuspended by pipetting 4-5 times with 1 ml Fixation buffer and fixed at 4°C for 16-24 hr. The fixed sample was then centrifuged at 850 x g for 5 min at room temperature. Supernatant was removed, and the pellet was resuspended by pipetting 4-5 times with 1 ml Quenching buffer. To ensure long-term stability at -80°C, 100 µl of Enhancer was added to the solution, followed by 275 µl 50% glycerol. A 50 µl aliquot was separated to assess RNA integrity and cell count. Both aliquot and main stock were frozen at -80°C. Sequencing of the samples was performed by the Center for Translational Genomics (CTG) at Lund University.

The data were pre-processed with CTG singleCellWorkflows – Nextflow pipeline (release 1.6.2 – containing CellRanger v7.1.0, FastQC v0.11.9, and MultiQC v1.16). Count matrixes with raw data were filtered with EmptyDrops (DropletUtils R package v1.16.0^66^), with FDR <= 0.01, to remove empty droplets, yielding approx. 5.7k – 7.6k cells per sample and ∼17.6k genes per sample. The data were further processed using R programming language (v4.2.2^67^) and the Seurat package (v4.3.0^68^). The data were split to two subsets – hCOs and hSOs – and integrated within each subset sample-wise. From now on, the two subsets were analysed separately, following standard Seurat processing pipeline (LogNormalize, ScaleData, and SCTransform). Quality control (QC) and filtering included identification of doublets with DoubletFinder (v2.0.4^69^, estimated doublet rate set to 5% according to 10x recommendation) and removal of cells with mitochondrial transcript abundance over 15% (keeping percent.mt < 15%) and less than 2000 genes (keeping nFeature > 2000). After these QC steps, the datasets contained 43,339 cells and 18,485 genes (hCOs), and 39,234 cells and 18,472 genes (hSOs). A small cluster of excitatory neurons, which was unique to a single sample (GD-iso at day 150), as well as several scattered clusters of low-quality cells, were excluded from the hSO dataset. Details can be found in Fig S8.

The data were visualized with UMAP and *FindNeighbors()* and *FindClusters()* functions were used to identify clusters. In hCOs, the integrated assay was used to better match the GD-iso DIV75 sample with the rest. In hSOs, the SCT assay was used for clustering. Cell cycle score was assigned to cells by *CellCycleScoring()* function. *FindAllMarkers(min.pct = 0.8, logfc.threshold = 0.65, only.pos = TRUE)* was used to identify cluster markers using the default Wilcoxon test. Monocle3 (v1.3.1^70–72^) was used for pseudotime trajectory construction, with “radial glia” and “early IN progenitors” clusters as the root cells in hCO and hSO subset, respectively. Differential abundance of populations was tested with scProportionTest (v0.0.0.9000) R package^73^.

The differential expression analysis (DEA) was performed using t-test in *FindMarkers(logfc.threshold = 0, min.pct = 0.1, test.use = “t”)* function. The significance threshold for differentially expressed genes (DEGs) was |log_2_FC| > 0.65 and p_adj_ < 0.05 with Bonferroni correction. Overrepresentation analysis of Gene Ontology terms in lists of DEGs was performed with clusterProfiler R package (v4.4.4^74^), function *compareCluster(fun = “enrichGO”, OrgDb = org.Hs.eg.db, keyType = “ALIAS”, ont = “ALL”, pvalueCutoff = 0.05)*. GSEA was performed with ranked gene lists (ranked by log2FC * -log10(padj)) using clusterProfiler function *gseGO(ont =“ALL”, keyType = “ALIAS”, minGSSize = 5, maxGSSize = 800, pvalueCutoff = 0.9, verbose = TRUE, OrgDb = org.Hs.eg.db, pAdjustMethod = “BH”, by = “fgsea”)*.

### Microelectrode arrays (MEA)

Day 150 or older hCOs and hSOs were individually transferred to MEA probes (60PedotMEA200/30iR-Au; Multichannel systems) with CO_2_ and temperature-equilibrated BrainPhys media using a wide-bore P1000 pipette tip, and a harp (HSG-MEA; ALA Scientific) was gently lowered to hold the organoid in place over the electrodes. The probe was then returned to a 37°C 5% CO_2_ 95% O_2_ incubator to stabilise for 30 min. Next, the probe was transferred to 37°C heated MEA headstage (MEA2100, Multichannel systems) and an environmental chamber with continuous perfusion of a 5% CO_2_ 95% O_2_ mix was secured over the MEA probe. After 10 min stabilisation, a 5 min baseline recording was made at 20 KHz. After baseline, half media was exchanged with pre-equilibriated BrainPhys with Picrotoxin (PTX) diluted to a final concentration of 100 µM followed by a 5 min equilibrium period and a 5 min recording period. Finally half media was exchanged with pre-equilibriated BrainPhys with PTX, NBQX and DL-APV diluted to final concentrations of 100, 5 and 100 µM respectively, followed by a 5 min equilibrium period then a 5 min recording period. Acquired raw traces were exported in .hdf5 format to allow their subsequent processing and spike/burst data extraction in Python. The code used for analysis is available on demand. Extracted values were imported into R Studio for analysis.

### Forward programmed tricultures

Generation of induced tricultures of induced neurons (iNs), interneurons (iGNs) and astrocytes (iA) was performed following minor modifications to previously published protocols^61, 75, 76^. Briefly, iPSCs at ∼80% confluency were dissociated with Accutase (Day -2) and 3 x 10^5^ (iNs), 3.5 x 10^5^ (iGNs) or 4 x 10^5^ cells were plated on Matrigel Growth Factor Reduced Basement Membrane Matrix (Corning) coated 6-well plates with mTeSR1 supplemented with 10 µM RI. The following day (day -1), media was replaced with fresh mTeSR1 media, and 1 µl of rtTA and Ngn2, Ascl1+Dlx2 or NFIB+SOX9 lentivirus were added to each well. Lentiviral vectors for FUW-M2-rtTA (reverse tetracycline-controlled transactivator, a gift from Rudolf Jaenisch (Addgene, #20342) and tetO.NGN2.puro, tetO.ASCL1.puro, tetO.DLX2.hygro a gift from Marius Wernig (Addgene, #52047, #97329, #97330) were prepared as previously described^77^. For all cell types on on day 0, media was replaced with fresh mTeSR1 media containing 2.5 µg / ml of doxycycline (Dox; Sigma-Aldrich), which was kept in the media throughout experiments. *For neurons:* From day 1-2, BrainPhys media (StemCell Technologies) supplemented with 0.5% N2 and 1% B27 supplement was used, and 48 h of 1.25 µg/ml puromycin ± 200 µg/mL hygromycin (hygromycin for interneurons only) selection was performed. From day 3-4 daily media changes continued without puromycin selection. From day 5-6, daily media exchanges were performed using BrainPhys supplemented with 0.5% N2, 1% B27, 10 ng/ml NT3 (Peprotech) and 10 ng / ml BDNF (Peprotech) ± 200 µg/mL hygromycin (hygromycin for interneurons only) selection. On day 5-6 for interneurons 5 µM AraC was supplemented into the media to remove highly mitotic cells.

For astrocytes: From day 1-2, expansion media, DMEM/F-12 supplemented with 10% FBS, 1% N2 supplement and 1% Glutamax was used, and 48 h of 1.25 µg/ml puromycin ± 200 µg/ml hygromycin was performed. From day 3-5 ± 200 µg/ml hygromycin selection was maintained whilst the media was transitioned from 3:1, 1:1 and 1:3 mixes of expansion media to FGF media, which was comprised of Neurobasal, 2% B27 supplement, 1% NEAA, 1% Glutamax, 1% FBS and supplemented with 8 ng/ml FGFb, 5 ng/ml CNTF and 10 ng/ml BMP4. On day 6 a full exchange of FGF media without hygromycin was performed.

At day 7 iNs, iGNs, and iAs were rinsed with DPBS without Ca and Mg, dissociated with Accutase, resuspended in 4 volumes DMEM:F12 supplemented with DNase I, pelleted at 300 x g and then resuspended in 1 ml (iNs) or 0.5 ml (iGNs, iAs) coculture media per original well. Clumps were then filtered via a 40 µm strainer and a cell count was performed before replating at a ratio of 6:3:2 iN:iGN:iA on the final culture surface. For IBIDI u-dish plates cultures were plated at 2.8 x 10^5^ total cells. From this point onwards media was exchanged every 3-4 days with a 1:1 mix of Brainphys supplemented with 2% B27, 1% N2, BDNF, NT3 and astrocyte maturation medium prepared as previously described.

### High density microelectrode arrays (HD-MEA)

Chimeric tricultures were prepared as previously described in “Forward programmed tricultures”, with minor modifications. MaxTwo 24-well plates were conditioned with 0.6 mL complete co-culture media for 48 h in a 5% CO2 37oC incubator at relative humidity. Wells were rinsed once with sterile ddH2O before drying, and 50 µl of 0.1 mg/ml PDL coating solution was spotted over the entire coating area before the plate was sealed with a Breathe-Easy membrane and incubated for 1 h at 37°C. After incubation, each well was rinsed three times with sterile ddH_2_O before drying for 1 h under a sterile airflow. 50 µl of 5 µg/ml RhLam521 (Biolamina) diluted in DPBS +/+ was added to the plating area and the plate was sealed with a breathe-easy membrane before storage overnight at 4°C. The following day, induced neurons, interneurons and astrocytes were resuspended as previously described, before being plated in a 6:3:2 ratio in 6.2 x 10^4^ cells in 50 µl coculture media supplemented with 10 µM Rho kinase inhibitor. A breathe easy membrane was applied, and the cells were incubated for 1 h before 0.6 ml coculture media supplemented with 10 µM Rho kinase media was added to the well. From this point on the same protocol was previously described was followed, with the addition of 1:100 Pencillin/Streptomycin to maintain sterility. From Day 14 – 54 plated chimeric tricultures on compatible MaxTwo 24-well plates were transferred to a MaxTwo HD-MEA system (MaxWell Biosystems) for recording with CO_2_ and temperature-equilibriation. The plate was allowed to acclimatise for 5 min before an activity scan (90 s, Checker Board) was performed to map the active electrodes. After an activity scan for the session was generated, it was utilised to select recording electrodes (Feature Maximization, Firing rate, 1020 Electrodes) for a 300 s baseline recording period. On Day 54, 100 µM Pictrotoxin or a DMSO control of the same volume was added to each well, allowed to stabilise for 10 min, and then another recording was acquired using the same activity scan as the prior recording.

Acquired raw traces were exported in .hdf5 format to allow their subsequent processing and spike/burst data extraction in Python. The code used for analysis is available on demand. Extracted values were imported into R Studio for analysis.

### Calcium imaging

Chimeric tricultures were prepared as previously described in “Forward programmed tricultures” and on Day 14 were transduced in 1 ml volume with 0.5 µl ssAAV-DJ/2-mCaMKIIa-jGCaMP8m-WPRE-bGHp(A) (v630-DJ; 9.0 x 10^12^ vg/ml) produced by the Viral Vector Facility (VVF) of the Neuroscience Center Zurich (ZNZ). On Day 20 cultures were transferred to BPIO media consisting of BrainPhys Imaging optimised medium (STEMCELL Technologies) supplemented with supplemented with 0.5% N2, 1% B27, 10 ng/ml NT3 (Peprotech) and 10 ng/ml BDNF (Peprotech). From Day 21-49 weekly recordings were performed consisting of a 3 min baseline recording window, followed by treatment with 10 µM Bicuculine, incubation for 5 min to restore CO^2^ and temperature, then a second 3 min treatment recording window. Cultures were then rinsed to remove residual bicuculine and reincubated with a 1:1 mix of fresh and pre-conditioned BPIO media. Recordings were performed on a custom Ultimeyes microscopy setup (Cairn GmBH). Images were acquired at 20 Hz using Micro-Manager to initalise acquisition via a CellCam Kikker camera, and illuminated with a CoolLED connected to a GFP filter. Recording analysis was performed using NetCal^78^ to extract calcium traces and analyse transient metrics.

## Statistical analysis

All experiments were performed with 3 biological replicates and results expressed as the mean ± SEM unless differently indicated in the figure legend, with statistical analysis using performed using Prism software with the tests indicated in the figure legends.

## ACKNOWLEDGMENTS

We would like to thank the Cell And Gene Therapy platform at Lund University for their support generating the iPSC lines; the Center for Translational Genomics at Lund University for their service performing the scRNAseq experiments; the Electrophysiology Unit at Lund Stem Cell Center for their help carrying out the organoid MEA recordings and analyses. We would also like to thank Prof. David Liu and Prof. Sabine Fuchs and their respective labs for advice on the design of pegRNAs for prime editing.

This work was supported by SSMF Stora Anslag (S20-0003), Hjärnfonden (FO2023-0391), Jeanssons Stiftelse (J2021-0018), Magnus Bergvalls Stiftelse, Segerfalk Stiftelse, Linnea and Josef Carlssons Stiftelse, Åhlén Stiftelse and Parkinson Schweiz to I.C. J.A.C., A.S.G and I.C. are supported by the University of Zurich Research Priority Program ITINERARE – Innovative Therapies in Rare Diseases. J.A.C. was supported by grants of the Royal Physiographic Society of Lund, Rut och Erik Hardebos Stiftelse for brain research, University of Zurich Postdocoral Fellowship Grant and Anna Mueller Grocholski-Stiftung; Z.M and L.V. were supported by the Czech Science Foundation (grants 24-11364S, 24-12028S) and by institutional support (RVO 86652036). J.B and U.E were supported by grants from Lund University and the Royal Physiographic Society of Lund;

## Notes

### Competing Interest Statement

The authors have declared no competing interest.

## REFERENCES

1. Platt, F.M., Boland, B. & van der Spoel, A.C. The cell biology of disease: lysosomal storage disorders: the cellular impact of lysosomal dysfunction. J Cell Biol 199, 723–734 (2012).

2. Saftig, P. & Klumperman, J. Lysosome biogenesis and lysosomal membrane proteins: trafficking meets function. Nat Rev Mol Cell Biol 10, 623–635 (2009).

3. Nalysnyk, L., Rotella, P., Simeone, J.C., Hamed, A. & Weinreb, N. Gaucher disease epidemiology and natural history: a comprehensive review of the literature. Hematology 22, 65–73 (2017).

4. Mazzulli, J.R. et al. Gaucher disease glucocerebrosidase and alpha-synuclein form a bidirectional pathogenic loop in synucleinopathies. Cell 146, 37–52 (2011).

5. Platt, F.M., d’Azzo, A., Davidson, B.L., Neufeld, E.F. & Tifft, C.J. Lysosomal storage diseases. Nat Rev Dis Primers 4, 27 (2018).

6. Aerts, J. et al. Glycosphingolipids and lysosomal storage disorders as illustrated by gaucher disease. Curr Opin Chem Biol 53, 204–215 (2019).

7. Stirnemann, J. et al. A Review of Gaucher Disease Pathophysiology, Clinical Presentation and Treatments. Int J Mol Sci 18 (2017).

8. Alcalay, R.N. et al. Age-Specific Parkinson Disease Risk in Gaucher Disease Type 1: Data From the ICGG Gaucher Registry. Neurology 106, e214986 (2026).

9. Reza, S., Ugorski, M. & Suchanski, J. Glucosylceramide and galactosylceramide, small glycosphingolipids with significant impact on health and disease. Glycobiology 31, 1416–1434 (2021).

10. Yu, R.K., Tsai, Y.T. & Ariga, T. Functional roles of gangliosides in neurodevelopment: an overview of recent advances. Neurochem Res 37, 1230–1244 (2012).

11. Giuffrida, G. et al. Glucosylsphingosine (Lyso-Gb1) as a reliable biomarker in Gaucher disease: a narrative review. Orphanet J Rare Dis 18, 27 (2023).

12. Farfel-Becker, T., Vitner, E.B. & Futerman, A.H. Animal models for Gaucher disease research. Dis Model Mech 4, 746–752 (2011).

13. Kim, E.Y., Hong, Y.B., Go, S.H., Lee, B. & Jung, S.C. Downregulation of neurotrophic factors in the brain of a mouse model of Gaucher disease; implications for neuronal loss in Gaucher disease. Exp Mol Med 38, 348–356 (2006).

14. Degl’Innocenti, E. & Dell’Anno, M.T. Human and mouse cortical astrocytes: a comparative view from development to morphological and functional characterization. Front Neuroanat 17, 1130729 (2023).

15. Siletti, K. et al. Transcriptomic diversity of cell types across the adult human brain. Science 382, eadd7046 (2023).

16. Bakken, T.E. et al. Comparative cellular analysis of motor cortex in human, marmoset and mouse. Nature 598, 111–119 (2021).

17. Hodge, R.D. et al. Conserved cell types with divergent features in human versus mouse cortex. Nature 573, 61–68 (2019).

18. van Hout, A.T.B., van Heukelum, S., Rushworth, M.F.S., Grandjean, J. & Mars, R.B. Comparing mouse and human cingulate cortex organization using functional connectivity. Brain Struct Funct 229, 1913–1925 (2024).

19. Birtele, M., Lancaster, M. & Quadrato, G. Modelling human brain development and disease with organoids. Nat Rev Mol Cell Biol 26, 389–412 (2025).

20. Tiscornia, G. et al. Neuronopathic Gaucher’s disease: induced pluripotent stem cells for disease modelling and testing chaperone activity of small compounds. Hum Mol Genet 22, 633–645 (2013).

21. Sun, Y. et al. Properties of neurons derived from induced pluripotent stem cells of Gaucher disease type 2 patient fibroblasts: potential role in neuropathology. PLoS One 10, e0118771 (2015).

22. Srikanth, M.P. & Feldman, R.A. Elevated Dkk1 Mediates Downregulation of the Canonical Wnt Pathway and Lysosomal Loss in an iPSC Model of Neuronopathic Gaucher Disease. Biomolecules 10 (2020).

23. Schondorf, D.C. et al. iPSC-derived neurons from GBA1-associated Parkinson’s disease patients show autophagic defects and impaired calcium homeostasis. Nat Commun 5, 4028 (2014).

24. Pornsukjantra, T. et al. An increase in ER stress and unfolded protein response in iPSCs-derived neuronal cells from neuronopathic Gaucher disease patients. Sci Rep 14, 9177 (2024).

25. Messelodi, D. et al. Neuronopathic Gaucher disease models reveal defects in cell growth promoted by Hippo pathway activation. Commun Biol 6, 431 (2023).

26. Brown, R.A. et al. mTOR hyperactivity mediates lysosomal dysfunction in Gaucher’s disease iPSC-neuronal cells. Dis Model Mech 12 (2019).

27. Baden, P. et al. Glucocerebrosidase is imported into mitochondria and preserves complex I integrity and energy metabolism. Nat Commun 14, 1930 (2023).

28. Awad, O. et al. Altered TFEB-mediated lysosomal biogenesis in Gaucher disease iPSC-derived neuronal cells. Hum Mol Genet 24, 5775–5788 (2015).

29. Aflaki, E. et al. A characterization of Gaucher iPS-derived astrocytes: Potential implications for Parkinson’s disease. Neurobiol Dis 134, 104647 (2020).

30. Rosety, I. et al. Impaired neuron differentiation in GBA-associated Parkinson’s disease is linked to cell cycle defects in organoids. NPJ Parkinsons Dis 9, 166 (2023).

31. Jo, J. et al. Lewy Body-like Inclusions in Human Midbrain Organoids Carrying Glucocerebrosidase and alpha-Synuclein Mutations. Ann Neurol 90, 490–505 (2021).

32. Frattini, E. et al. Lewy pathology formation in patient-derived GBA1 Parkinson’s disease midbrain organoids. Brain 148, 1242–1257 (2025).

33. Lin, Y. et al. Patient-specific midbrain organoids with CRISPR correction recapitulate neuronopathic Gaucher disease phenotypes and enable evaluation of novel therapies. Elife 15 (2026).

34. Zetterdahl, O.G. et al. Generation of iPSC lines with tagged alpha-synuclein for visualization of endogenous protein in human cellular models of neurodegenerative disorders. eNeuro (2025).

35. Ludwig, T.E. et al. ISSCR standards for the use of human stem cells in basic research. Stem Cell Reports 18, 1744–1752 (2023).

36. Anzalone, A.V. et al. Search-and-replace genome editing without double-strand breaks or donor DNA. Nature 576, 149–157 (2019).

37. Sloan, S.A. et al. Human Astrocyte Maturation Captured in 3D Cerebral Cortical Spheroids Derived from Pluripotent Stem Cells. Neuron 95, 779–790 e776 (2017).

38. Palmano, K., Rowan, A., Guillermo, R., Guan, J. & McJarrow, P. The role of gangliosides in neurodevelopment. Nutrients 7, 3891–3913 (2015).

39. Sipione, S., Monyror, J., Galleguillos, D., Steinberg, N. & Kadam, V. Gangliosides in the Brain: Physiology, Pathophysiology and Therapeutic Applications. Front Neurosci 14, 572965 (2020).

40. Zhang, K. et al. Imbalance of Excitatory/Inhibitory Neuron Differentiation in Neurodevelopmental Disorders with an NR2F1 Point Mutation. Cell Rep 31, 107521 (2020).

41. Furderer, M.L., Hertz, E., Lopez, G.J. & Sidransky, E. Neuropathological Features of Gaucher Disease and Gaucher Disease with Parkinsonism. Int J Mol Sci 23 (2022).

42. Chida, R., Shimura, M., Ishida, Y., Suganami, Y. & Yamanaka, G. Perinatal lethal Gaucher disease: A case report and review of literature. Brain Dev 45, 134–139 (2023).

43. Gornati, R. et al. Glycolipid analysis of different tissues and cerebrospinal fluid in type II Gaucher disease. J Inherit Metab Dis 25, 47–55 (2002).

44. Blumenreich, S. et al. Elevation of gangliosides in four brain regions from Parkinson’s disease patients with a GBA mutation. NPJ Parkinsons Dis 8, 99 (2022).

45. Ceni, C. et al. Identification of GM1-Ganglioside Secondary Accumulation in Fibroblasts from Neuropathic Gaucher Patients and Effect of a Trivalent Trihydroxypiperidine Iminosugar Compound on Its Storage Reduction. Molecules 29 (2024).

46. Plotegher, N. & Duchen, M.R. Mitochondrial Dysfunction and Neurodegeneration in Lysosomal Storage Disorders. Trends Mol Med 23, 116–134 (2017).

47. Kann, O., Papageorgiou, I.E. & Draguhn, A. Highly energized inhibitory interneurons are a central element for information processing in cortical networks. J Cereb Blood Flow Metab 34, 1270–1282 (2014).

48. Liu, Y. et al. Constraint-based modelling of metabolic dysregulation in Gaucher disease: mitochondrial dysfunction and disrupted cholesterol homeostasis. Orphanet J Rare Dis 21 (2026).

49. Garcia-Sanz, P., J, M.F.G.A. & Moratalla, R. The Role of Cholesterol in alpha-Synuclein and Lewy Body Pathology in GBA1 Parkinson’s Disease. Mov Disord 36, 1070–1085 (2021).

50. Sezgin, E., Levental, I., Mayor, S. & Eggeling, C. The mystery of membrane organization: composition, regulation and roles of lipid rafts. Nat Rev Mol Cell Biol 18, 361–374 (2017).

51. Duchemin, A.M., Ren, Q., Neff, N.H. & Hadjiconstantinou, M. GM1-induced activation of phosphatidylinositol 3-kinase: involvement of Trk receptors. J Neurochem 104, 1466–1477 (2008).

52. Caracci, M.O., Fuentealba, L.M. & Marzolo, M.P. Golgi Complex Dynamics and Its Implication in Prevalent Neurological Disorders. Front Cell Dev Biol 7, 75 (2019).

53. Subkhangulova, A. & Mikhaylova, M. The Golgi apparatus: adaptations to neuronal shape and functions. EMBO J 45, 358–373 (2026).

54. Sano, R. et al. GM1-ganglioside accumulation at the mitochondria-associated ER membranes links ER stress to Ca(2+)-dependent mitochondrial apoptosis. Mol Cell 36, 500–511 (2009).

55. Weesner, J.A. et al. Altered GM1 catabolism affects NMDAR-mediated Ca(2+) signaling at ER-PM junctions and increases synaptic spine formation in a GM1-gangliosidosis model. Cell Rep 43, 114117 (2024).

56. Yang, X. et al. Electroencephalogram and phenotype patterns in neuronopathic Gaucher disease patients - ten years of experience in a single center. Acta Epileptol 6, 30 (2024).

57. van van Hugte, E.J.H., Schubert, D. & Nadif Kasri, N. Excitatory/inhibitory balance in epilepsies and neurodevelopmental disorders: Depolarizing gamma-aminobutyric acid as a common mechanism. Epilepsia 64, 1975–1990 (2023).

58. Canals, I. et al. Astrocyte dysfunction and neuronal network hyperactivity in a CRISPR engineered pluripotent stem cell model of frontotemporal dementia. Brain Commun 5, fcad158 (2023).

59. Hsu, J.Y. et al. PrimeDesign software for rapid and simplified design of prime editing guide RNAs. Nat Commun 12, 1034 (2021).

60. Hsu, P.D. et al. DNA targeting specificity of RNA-guided Cas9 nucleases. Nat Biotechnol 31, 827–832 (2013).

61. Zhang, Y. et al. Rapid single-step induction of functional neurons from human pluripotent stem cells. Neuron 78, 785–798 (2013).

62. Quist, E., Ahlenius, H. & Canals, I. Transcription Factor Programming of Human Pluripotent Stem Cells to Functionally Mature Astrocytes for Monocultures and Cocultures with Neurons. Methods Mol Biol 2352, 133–148 (2021).

63. Chabas, A. et al. Neuronopathic and non-neuronopathic presentation of Gaucher disease in patients with the third most common mutation (D409H) in Spain. J Inherit Metab Dis 19, 798–800 (1996).

64. Yoon, S.J. et al. Reliability of human cortical organoid generation. Nat Methods 16, 75–78 (2019).

65. Birey, F. et al. Assembly of functionally integrated human forebrain spheroids. Nature 545, 54–59 (2017).

66. Lun, A.T.L. et al. EmptyDrops: distinguishing cells from empty droplets in droplet-based single-cell RNA sequencing data. Genome Biol 20, 63 (2019).

67. R Core Team (R Foundation for Statistical Computing, Vienna, Austria; 2022).

68. Hao, Y. et al. Integrated analysis of multimodal single-cell data. Cell 184, 3573–3587 e3529 (2021).

69. McGinnis, C.S., Murrow, L.M. & Gartner, Z.J. DoubletFinder: Doublet Detection in Single-Cell RNA Sequencing Data Using Artificial Nearest Neighbors. Cell Syst 8, 329–337 e324 (2019).

70. Trapnell, C. et al. The dynamics and regulators of cell fate decisions are revealed by pseudotemporal ordering of single cells. Nat Biotechnol 32, 381–386 (2014).

71. Qiu, X. et al. Reversed graph embedding resolves complex single-cell trajectories. Nat Methods 14, 979–982 (2017).

72. Cao, J. et al. The single-cell transcriptional landscape of mammalian organogenesis. Nature 566, 496–502 (2019).

73. Miller, S.A. et al. LSD1 and Aberrant DNA Methylation Mediate Persistence of Enteroendocrine Progenitors That Support BRAF-Mutant Colorectal Cancer. Cancer Res 81, 3791–3805 (2021).

74. Wu, T. et al. clusterProfiler 4.0: A universal enrichment tool for interpreting omics data. Innovation (Camb) 2, 100141 (2021).

75. Yang, N. et al. Generation of pure GABAergic neurons by transcription factor programming. Nat Methods 14, 621–628 (2017).

76. Canals, I. et al. Rapid and efficient induction of functional astrocytes from human pluripotent stem cells. Nat Methods 15, 693–696 (2018).

77. Ansorge, S. et al. Development of a scalable process for high-yield lentiviral vector production by transient transfection of HEK293 suspension cultures. J Gene Med 11, 868–876 (2009).

78. Javier G. Orlandi, S.F.-G., Andrea Comella-Bolla, Mercè Masana, Gerardo García-Díaz Barriga, Mohammad Yaghoubi, Alexander Kipp, Josep M. Canals, Michael A. Colicos, Jörn Davidsen, Jordi Alberch, Jordi Soriano, Vol. Neuroscience (2017).

