## Supplementary figures and tables for "Isogenic forebrain organoids uncover early neurodevelopmental alterations and imbalances in neuronal function leading to hyperexcitation in Gaucher disease"

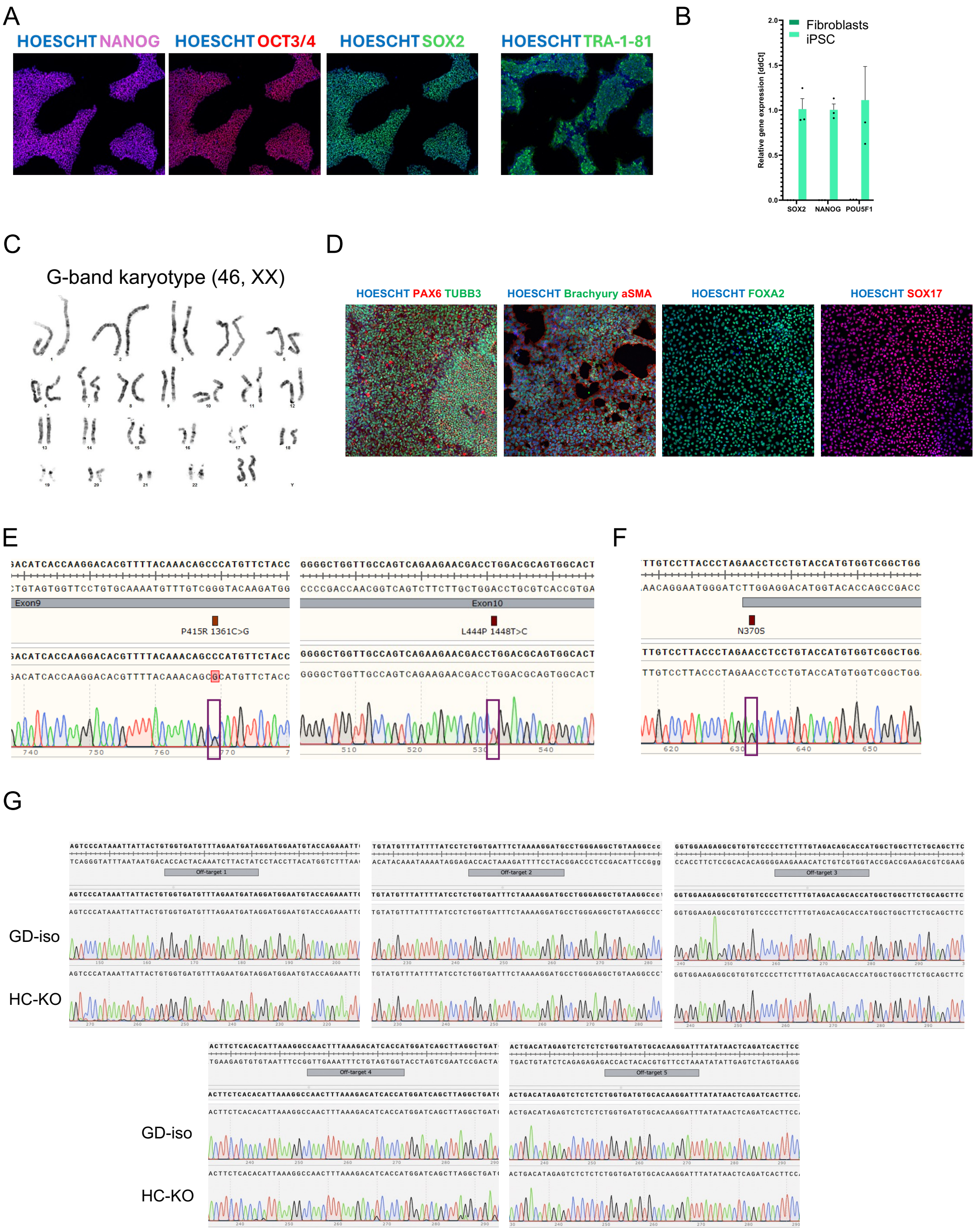


**Figure S1. Generation and characterization of a GD patient iPSC and isogenic lines with genome editing. A)** Representative images of immunocytochemistry assays for the markers of undifferentiated state NANOG, OCT3/4, SOX2 and TRA-1-81. **B)** Real-time quantitative PCR for markers of undifferentiated state SOX2, NANOG and POU5F1 normalised by GAPDH expression. **C)** Representative image of the G-band karyotyping for the GD iPSC line. **D)** Representative images of immunocytochemistry for markers of ectoderm (TUBB3, PAX6), mesoderm (SMA), and endoderm (FOXA2 and SOX17) after trilineage differentiation of the GD line. **E)** Chromatograms showing the presence of the mutations P454R (P415R) and L483P (L444P) in the GD line. **F)** Chromatogram showing the presence of the mutation N409S (historically known as N370S) in the HC line. **G)** Chromatograms showing no nucleotide changes for both the GD-iso and the HC-KO lines at the top-5 off-target sites for the sgRNA4.

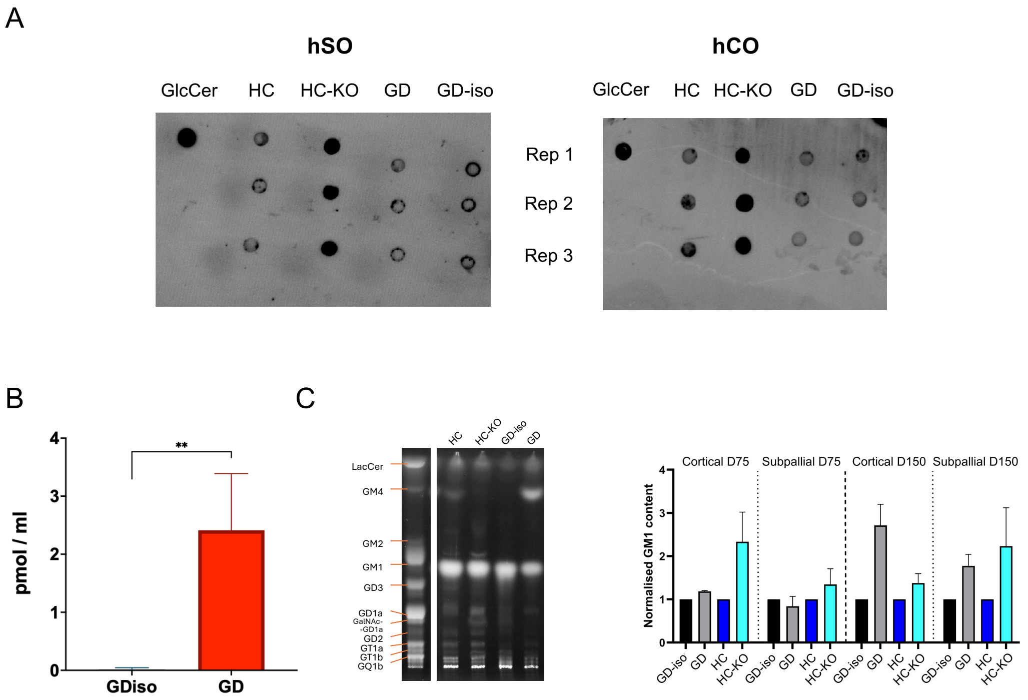


**Figure S2. Gaucher disease forebrain organoids display lipid accumulation. A)** Unprocessed dot blot images for GlcCer for hSOs and hCOs generated from all iPSC lines. **B)** GlcSph quantification in pmol per ml of sample in hCO at DIV75 comparing GD to GD-iso. Bar graphs represent the mean ± s.e.m. of 3 independent experiments. **p<0.01, one-tailed paired t-test. **C)** Representative image of a thin layer chromatography for hCOs generated from one replicate of all lines with a ganglioside control in the left lane; and bar graph showing the quantification of the normalised $GM_1$ bands from the thin layer chromatographies comparing GD-iso to GD and HC-KO to HC. Bar graphs represent the mean ± s.e.m. of 3 independent experiments.

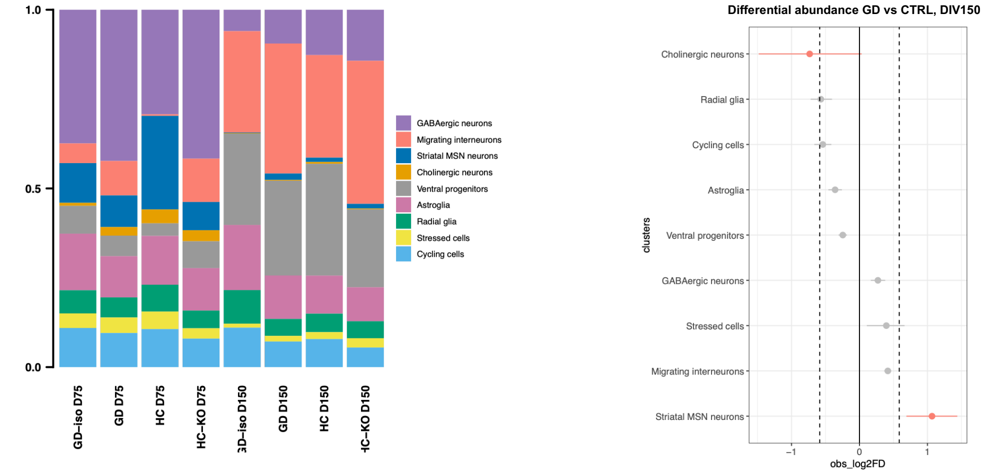


C

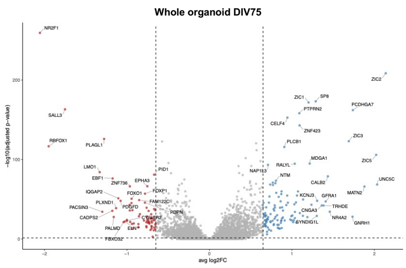


D

| Group | GD | CTRL | Up | Down |
|---|---|---|---|---|
| Whole organoid | 8927 | 7521 | 114 | 70 |
| GABAergic neurons | 3748 | 2512 | 89 | 43 |
| Migrating interneurons | 973 | 227 | 89 | 29 |
| Striatal MSN neurons | 747 | 1393 | 67 | 19 |
| Cholinergic neurons | 245 | 175 | 7 | 7 |
| Ventral progenitors | 593 | 428 | 51 | 86 |
| Astroglia | 1046 | 1111 | 102 | 47 |
| Radial glia | 471 | 526 | 18 | 22 |
| Stressed cells | 322 | 337 | 67 | 2 |
| Cycling cells | 782 | 812 | 54 | 41 |

E

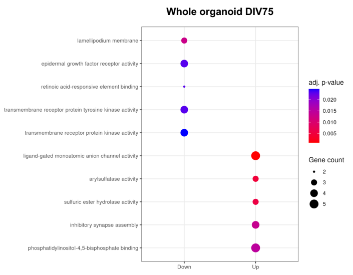


F

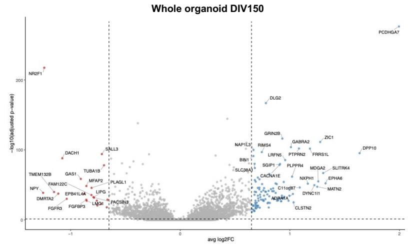


G

| Group | GD | CTRL | Up | Down |
|---|---|---|---|---|
| Whole organoid | 12368 | 10418 | 106 | 22 |
| GABAergic neurons | 1454 | 1015 | 32 | 8 |
| Migrating interneurons | 4714 | 2974 | 184 | 14 |
| Striatal MSN neurons | 194 | 78 | 3 | 3 |
| Cholinergic neurons | 25 | 35 | 0 | 0 |
| Ventral progenitors | 3004 | 3000 | 19 | 3 |
| Astroglia | 1348 | 1456 | 92 | 61 |
| Radial glia | 586 | 732 | 15 | 14 |
| Stressed cells | 256 | 164 | 2 | 1 |
| Cycling cells | 787 | 964 | 39 | 28 |

H

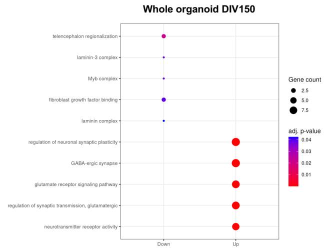


**Figure S3. Single-cell RNA sequencing of hSO display neurodevelopmental alterations. A)** Proportional bar plots showing the abundance of cells for each cluster in each sample and timepoint. **B)** Differential abundance of populations comparing GD vs CTRL at DIV150. **C)** Volcano plot displaying upregulated (up, blue) and downregulated (down, red) in whole hSO comparing GD vs CTRL at DIV75. **D)** Summary table displaying number of cells for each genotype and number of upregulated (up) and downregulated (down) genes for whole organoids and for each cluster at DIV75. **E)** Overrepresentation analysis results showing top five downregulated (down) and upregulated (up) GO terms distinguishing whole hSO from GD vs CTRL at DIV75. **F)** Volcano plot displaying upregulated (up, blue) and downregulated (down, red) in whole hSO comparing GD vs CTRL at DIV150. **G)** Summary table displaying number of cells for each genotype and number of upregulated (up) and downregulated (down) genes for whole organoids and for each cluster at DIV150. **H)** Overrepresentation analysis results showing top five downregulated (down) and upregulated (up) GO terms distinguishing whole hSO from GD vs CTRL at DIV150.

A

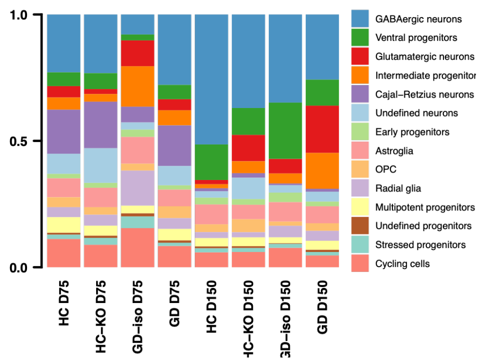


B

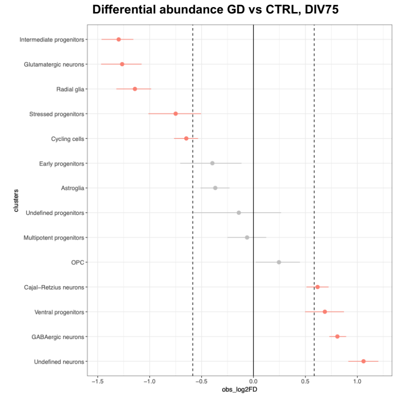


C

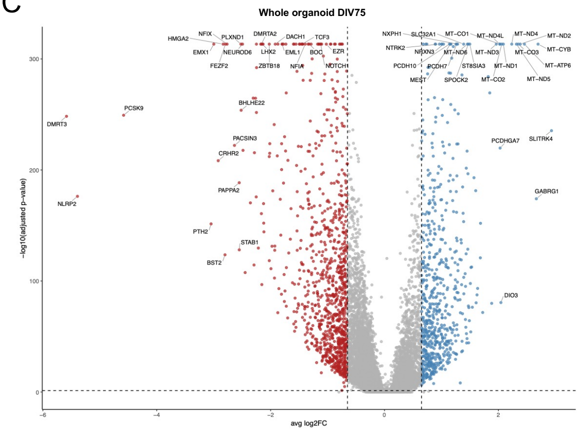


D

| Group | GD | CTRL | Up | Down |
|---|---|---|---|---|
| Whole organoid | 8714 | 10604 | 659 | 1008 |
| GABAergic neurons | 2227 | 1548 | 89 | 549 |
| Ventral progenitors | 520 | 393 | 52 | 190 |
| Glutamatergic neurons | 279 | 816 | 326 | 570 |
| Intermediate progenitors | 389 | 1164 | 596 | 954 |
| Cajal-Retzius neurons | 1505 | 1194 | 192 | 766 |
| Undefined neurons | 925 | 540 | 124 | 388 |
| Early progenitors | 158 | 253 | 191 | 579 |
| Astroglia | 624 | 980 | 632 | 1231 |
| OPC | 336 | 345 | 108 | 110 |
| Radial glia | 372 | 999 | 502 | 882 |
| Multipotent progenitors | 374 | 475 | 96 | 178 |
| Undefined progenitors | 76 | 102 | 36 | 55 |
| Stressed progenitors | 174 | 356 | 258 | 865 |
| Cycling cells | 755 | 1439 | 484 | 726 |

E

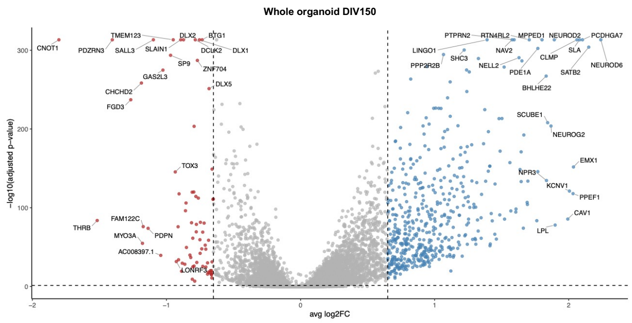


F

| Group | GD | CTRL | Up | Down |
|---|---|---|---|---|
| Whole organoid | 11608 | 12413 | 498 | 72 |
| GABAergic neurons | 3719 | 5356 | 203 | 39 |
| Ventral progenitors | 1216 | 2252 | 455 | 50 |
| Glutamatergic neurons | 1637 | 465 | 49 | 25 |
| Intermediate progenitors | 1031 | 341 | 37 | 12 |
| Cajal-Retzius neurons | 177 | 118 | 13 | 9 |
| Undefined neurons | 746 | 338 | 55 | 42 |
| Early progenitors | 240 | 399 | 73 | 18 |
| Astroglia | 718 | 969 | 163 | 105 |
| OPC | 488 | 295 | 16 | 8 |
| Radial glia | 330 | 425 | 114 | 71 |
| Multipotent progenitors | 413 | 355 | 93 | 95 |
| Undefined progenitors | 88 | 57 | 0 | 0 |
| Stressed progenitors | 174 | 199 | 31 | 52 |
| Cycling cells | 631 | 844 | 412 | 46 |

**Figure S4. Single-cell RNA sequencing of hCO reveal alterations in normal brain development. A)** Proportional bar plots showing the abundance of cells for each cluster in each sample and timepoint. **B)** Differential abundance of populations comparing GD vs CTRL at DIV75. **C)** Volcano plot displaying upregulated (up, blue) and downregulated (down, red) in whole hCO comparing GD vs CTRL at DIV75. **D)** Summary table displaying number of cells for each genotype and number of upregulated (up) and downregulated (down) genes for whole hCO and for each cluster at DIV75. **E)** Volcano plot displaying upregulated (up, blue) and downregulated (down, red) in whole hCO comparing GD vs CTRL at DIV150. **F)** Summary table displaying number of cells for each genotype and number of upregulated (up) and downregulated (down) genes for whole hCO and for each cluster at DIV150.

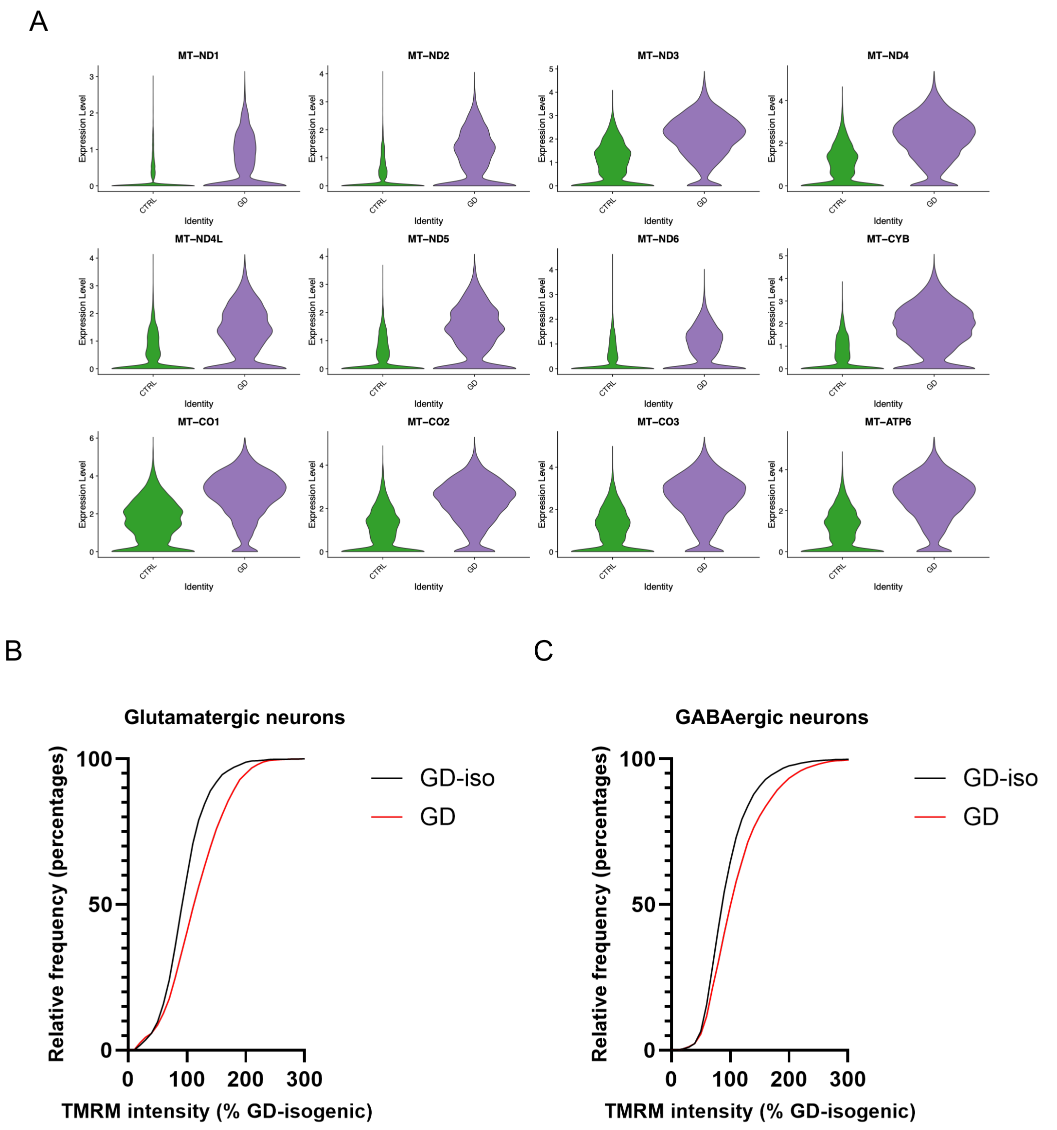


**Figure S5. Mitochondrial changes in hCO, glutamatergic neurons and GABAergic neurons. A)** Violin plots showing expression levels of selected mitochondrial genes in hCOs at DIV75 comparing GD (GD and HC-KO) with CTRL (GD-iso and HC) organoids. **B)** Relative frequency graph displaying fluorescence intensity distribution for glutamatergic neurons generated from GD and GD-iso iPSC. **C)** Relative frequency graph displaying fluorescence intensity distribution for GABAergic neurons generated from GD and GD-iso iPSC.

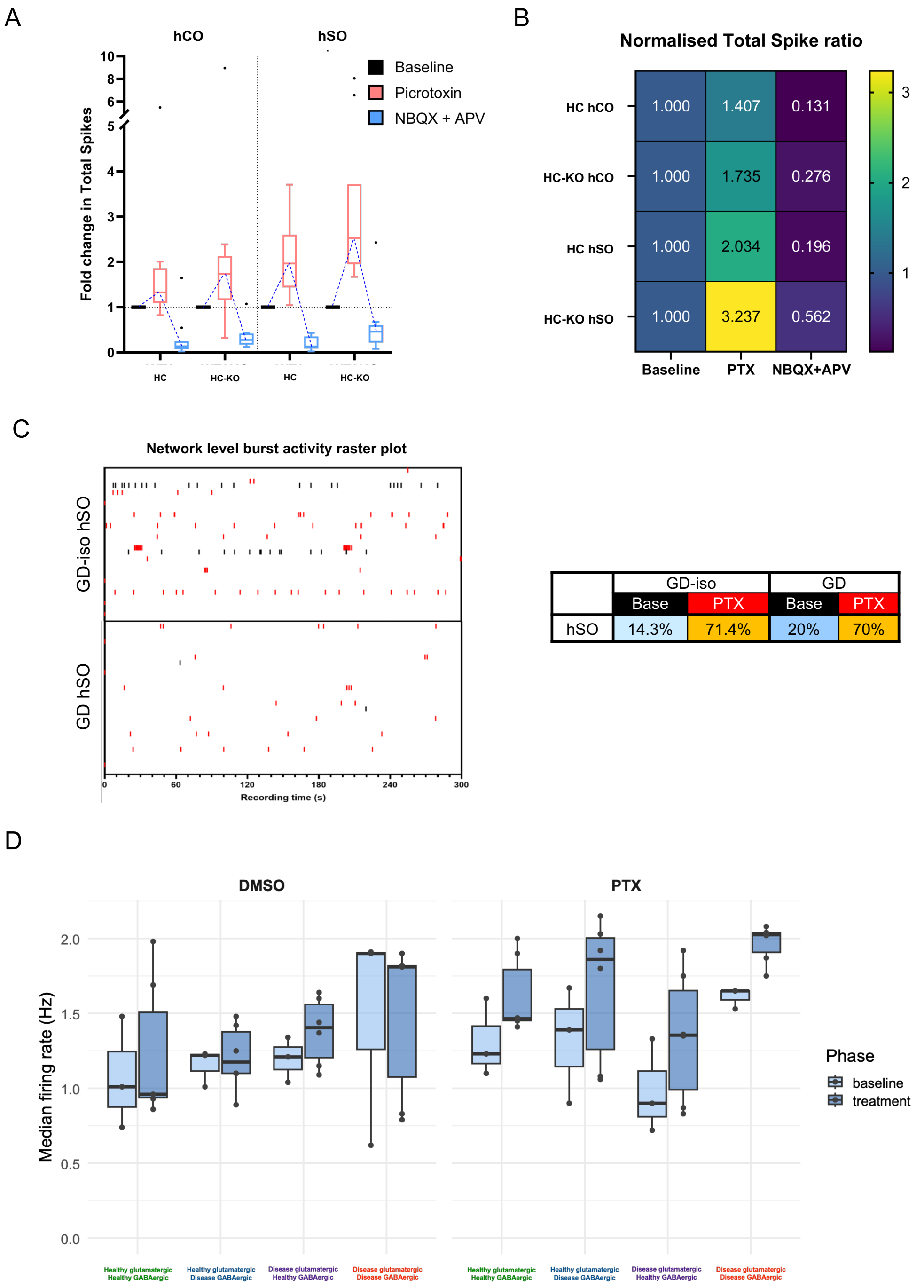


**Figure S6. MEA recordings show a hyperexcitability phenotype in Gaucher disease organoids and tricultures. A)** Fold-change in total spike number of hCO and hSO at baseline (black), after PTX addition (red) and after NBQX and APV addition (blue), for HC and HC-KO organoids at DIV150. **B)** Normalised total spike ratio for each type of organoid (hCO and hSO) generated from the HC and HC-KO lines comparing PTX or NBQX + APV treatment to the baseline recordings (bottom graph). **C)** Raster plot showing burst activity detection from individual hCOs from GD-iso and GD lines at baseline (black) or after PTX addition (red) for a 5 min recording, with the percentage of active electrodes illustrated in the table for each condition. **D)** Bar graph showing changes in median firing rate in 2D cultures of different combinations of healthy and disease glutamatergic and GABAergic neurons before and after vehicle (DMSO) or PTX addition.

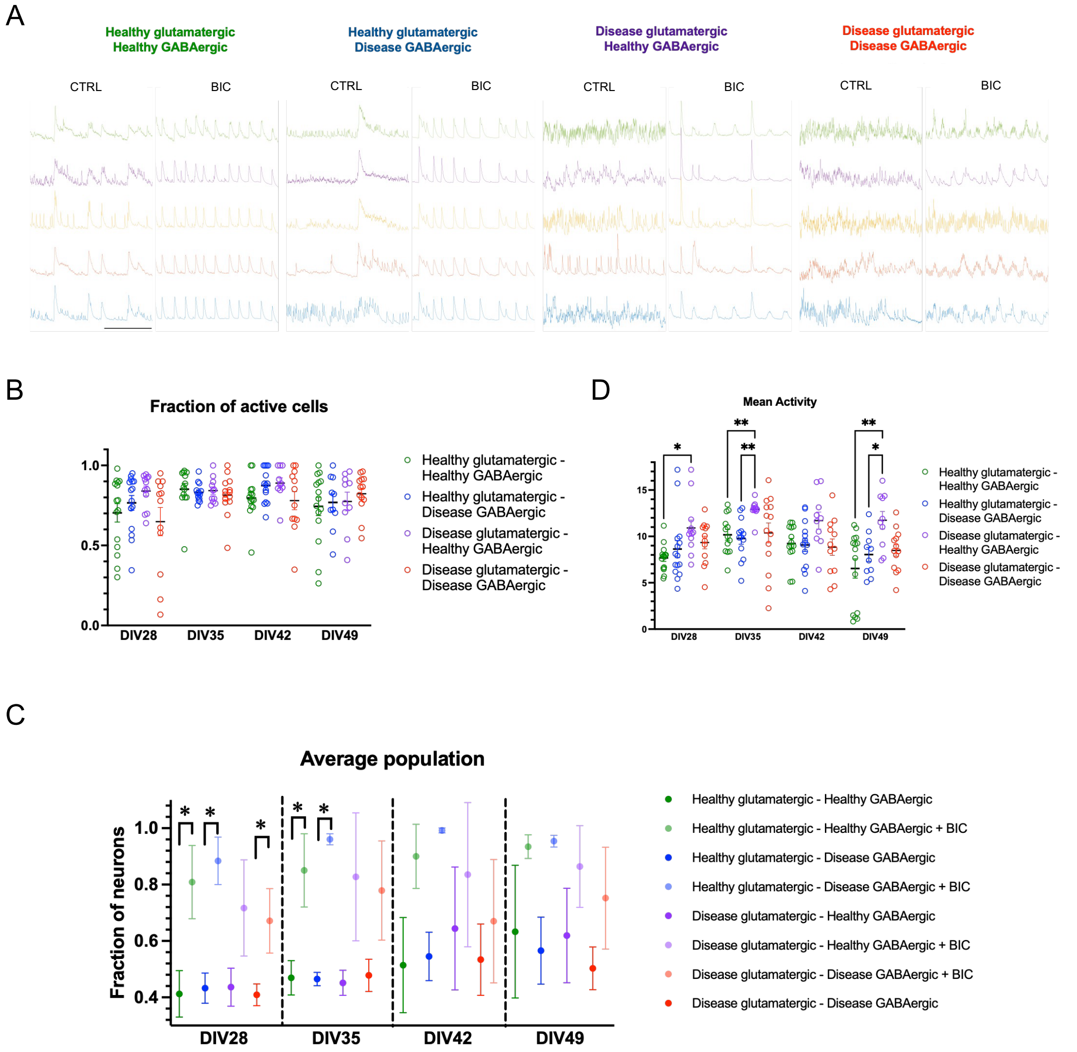


**Figure S7. Calcium imaging of CAMKIIa-GCaMP8m in chimeric forward programmed tricultures. A)** Representative traces from 5 cells of each condition at DIV49 before (CTRL) and after (BIC) bicuculline addition. **B)** Graph displaying the fraction of active cells over time (from DIV28 to DIV49) for each cell culture combination. **C)** Graph displaying mean activity of neurons over time (from DIV28 to DIV49) for each cell culture combination. **D)** Graph showing the average population represented as the fraction of neurons participating in collective events over time (from DIV28 to DIV49) before and after (BIC) bicuculline addition for each culture combination.

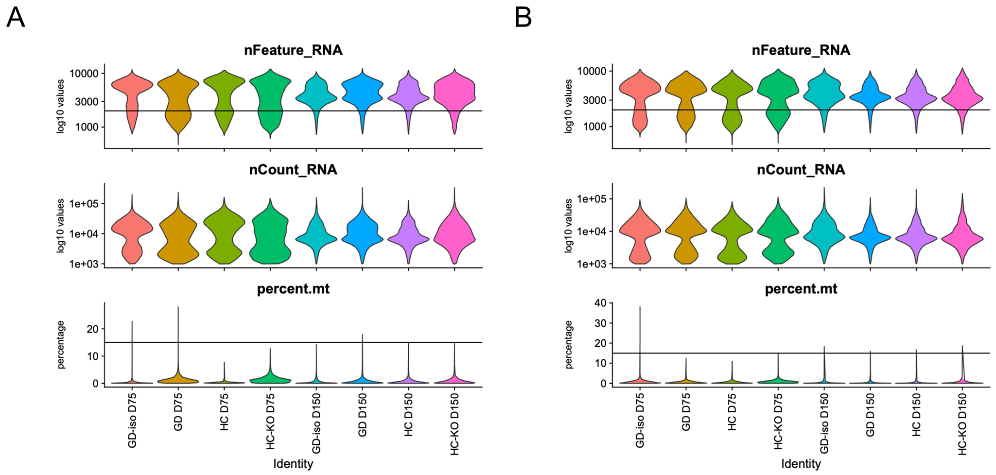


C

| Subset | SampleID | # of cells before QC, after EmptyDrops | # of cells after QC | # of non-zero genes before QC | # of non-zero genes after QC | median genes per cell (after QC) | median UMIs per cell (after QC) |
|---|---|---|---|---|---|---|---|
| hCS | GD-iso D75 | 7540 | 5873 | 17736 | 17456 | 5593 | 14287 |
| | GD D75 | 6738 | 4397 | 17800 | 17628 | 5030 | 12245 |
| | HC D75 | 6642 | 4731 | 17758 | 17642 | 5981 | 17176 |
| | HC-KO D75 | 6424 | 4317 | 17840 | 17630 | 4993 | 12106 |
| | GD-iso D150 | 6927 | 6199 | 17401 | 17204 | 3865 | 7957 |
| | GD D150 | 5759 | 5133 | 17430 | 17207 | 4533 | 10233 |
| | HC D150 | 7056 | 6214 | 17301 | 17247 | 3844 | 7770 |
| | HC-KO D150 | 7514 | 6475 | 17941 | 17769 | 4524 | 10036 |
| hVS | GD-iso D75 | 5989 | 3826 | 17602 | 17373 | 5034.5 | 12265.5 |
| | GD D75 | 7265 | 4419 | 17497 | 17294 | 5077 | 12762 |
| | HC D75 | 6311 | 3695 | 17665 | 17274 | 4967 | 12505 |
| | HC-KO D75 | 7548 | 4508 | 17937 | 17736 | 4919 | 11972.5 |
| | GD-iso D150 | 7651 | 4575 | 17514 | 16948 | 3668 | 7191 |
| | GD D150 | 7323 | 6474 | 17341 | 16996 | 3590 | 6953.5 |
| | HC D150 | 6700 | 5843 | 17264 | 17009 | 3427 | 6653 |
| | HC-KO D150 | 6866 | 5894 | 17624 | 17441 | 3480.5 | 6619 |

**Figure S8. Single-cell quality control and filtering of the data. A)** Quality control for hCO. Lines in nFeature_RNA (number of genes) and percent.mt (percentage of mitochondrial transcripts) indicate threshold values for filtering steps. nCount_RNA represents number of transcripts. **B)** Quality control for hSO. Lines in nFeature_RNA (number of genes) and percent.mt (percentage of mitochondrial transcripts) indicate threshold values for filtering steps. nCount_RNA represents number of transcripts. **C)** Table summarizing sequencing details for each sample. QC = quality control. UMIs = unique molecular identifiers.

**Table S1. Cell lines included in this study**

| Line | Coriell ID | Sex | Age | Genotype | Phenotype |
|---|---|---|---|---|---|
| **HC** | GM08398 | M | 8 y | WT / N370S | Healthy (carrier) |
| **HC-KO** | GM08398-derived | M | 8 y | K447fsX21 / F436_T446delinsS | *GBA1* knockout |
| **GD** | GM01260 | F | 11 mo | P415R / L444P | GD type II |
| **GD-iso** | GM01260-derived | F | 11 mo | WT / WT | Healthy |

**Table S2. Primers for amplifying *GBA1* exons**

| Exon | Forward strand primer | Reverse strand primer |
|---|---|---|
| Exon 1-2 | *5'-ATCCTCTGGGATTTAGGAGC-3'* | *5'-AGAAGGGAGGCTCTGTGCTA-3'* |
| Exon 3 | *5'-CCGTGTTCAGTCTCTCCTAG-3'* | *5'-GGAGGACCCAGCCTGGCCCA-3'* |
| Exon 4 | *5'-TGGGTACTGATACCCTTATT-3'* | *5'-TCAATGGCTCTATGTCATCT-3'* |
| Exon 5 | *5'-ACCCAGGAGCCCAAGTTCCC-3'* | *5'-CCTCAGGGCCTGAAAAAGCT-3'* |
| Exon 6 | *5'-CTCTGGGTGCTTCTCTCTTC-3'* | *5'-ACAGATCAGCATGGCTAAAT-3'* |
| Exon 7a | *5'-CTCGGCTTCCCAAAGTGC-3'* | *5'-CTAGGTCACGGGCAATGAAG-3'* |
| Exon 7b | *5'-TGGGCTTCACCCCTGAACAT-3'* | *5'-ATAGTTGGGTAGAGAAATCG-3'* |
| Exon 8-10 | *5'-TGTGCAAGGTCCAGGATCAG-3'* | *5'-TTCAGCCCACTTCCCAGACC-3'* |
| Exon 9-11 | *5'-CTGGAACCTTGCCCTGAAC-3'* | *5'-CTCTTTAGTCACAGACAGCG-3'* |

**Table S3. Thermocycler settings for amplifying *GBA1* exons**

| Exon | Initial denaturation | 32 – 34x cycles | | | Final extension |
|---|---|---|---|---|---|
| | | Denaturation | Annealing | Extension | |
| Exon 1-2 | 98°C 30s | 98°C 10s | 67.5°C 30s | 72°C 30s | 72°C 5m |
| Exon 3 | | | 61.9°C 30s | 72°C 15s | |
| Exon 4 | | | 62.5°C 30s | 72°C 15s | |
| Exon 5 | | | 70°C 30s | 72°C 15s | |
| Exon 6 | | | 65°C 30s | 72°C 15s | |
| Exon 7a | | | 60.7°C 30s | 72°C 15s | |
| Exon 7b | | | 61.5°C 30s | 72°C 15s | |
| Exon 8-10 | | | 68.5°C 30s | 72°C 45s | |
| Exon 9-11 | | | 65°C 30s | 72°C 30s | |

**Table S4. Antibodies used in this study**

| Target | Species | Dilution factor | Catalog ID | Manufacturer |
|---|---|---|---|---|
| NANOG | Rabbit | 1:100 | ab21624 | Abcam |
| OCT 3/4 (C10) | Mouse | 1:100 | sc-5279 | Santa Cruz Biotechnology |
| SOX2 | Goat | 1:100 | AF2018 | R&D Systems |
| SOX17 | Goat | 1:100 | AF1924 | R&D Systems |
| TRA1-81 M | Mouse | 1:200 | MAB4381 | Millipore |
| hHNF-3b / FOXA2 | Goat | 1:100 | AF2400 | R&D Systems |
| Brachyury | Goat | 1:100 | AF2085 | R&D Systems |
| α-fetoprotein | Rabbit | 1:100 | A0008 | Dako |
| α-Smooth Muscle Actin | Mouse | 1:500 | A5228 | Sigma-Aldrich |
| ßIII-tubulin | Mouse | 1:1000 | PRB-435P | BioLegend |
| ßIII-tubulin | Mouse | 1:500 | T8660 | Merck |
| PAX6 | Rabbit | 1:100 | sc-32766 | Santa Cruz Technology |
| PAX6 | Rabbit | 1:300 | 901301 | BioLegend |
| NeuN | Mouse | 1:1000 | ab104224 | Abcam |
| Alexa fluor 488 anti-Mouse (IgM) | Donkey | 1:500 | 715-545-140 | Jackson Immunoresearch |
| Alexa Fluor 568 anti-Rabbit | Donkey | 1:500 | A-10042 | Thermo Fisher Scientific |

| | | | | |
|---|---|---|---|---|
| Alexa Fluor 488 anti-Mouse (IgG) | Donkey | 1:500 | A-21202 | Thermo Fisher Scientific |
| Alexa Fluor 568 anti-Goat | Donkey | 1:500 | A-11057 | Thermo Fisher Scientific |

**Table S5. TaqMan assays for RT-PCR used in this study**

| Assay ID | Gene symbol | Lot number |
|---|---|---|
| Hs03044281_g1 | YWHAZ; reference gene | 2071687 |
| Hs00742896_s1 | POU5F1 | 2193241 |
| Hs02387400_g1 | NANOG | 2207564 |
| Hs01053049_s1 | SOX2 | 2199578 |

**Table S6. CRISPR/Cas9 sgRNA and ssODN used in this study.**

| Name | Sequence |
|---|---|
| P415R sgRNA-4 | 5'-CATCATTGTAGACATCACCA AGG-3' |
| P415R ssODN | 5'-T*T*GCTGAAGTGGCCAAGGTGGTAGAACATGGGCTGTTTGT AAAACGTGTCTTTGGTGATGTCTACAATGATGGGACTGT*C*G-3' |

**Table S7. L444P PE3b oligonucleotide sequences**

| | |
|---|---|
| **pegRNA Spacer sequences** | |
| Top strand | 5'-caccGCATCAGTGCCACTGCGTCCgtttt-3' |
| Bottom strand | 5'-ctctaaaacGGACGCAGTGGCACTGATGC-3' |
| **pegRNA Extension sequences** | |
| Top strand | 5'-gtgcAGAACgatCtGGACGCAGTGGCACTG-3' |
| Bottom strand | 5'-aaaaCAGTGCCACTGCGTCCaGatcGTTCT-3' |
| **ngRNA Complement sequences** | |
| Top strand | 5'-caccGTGCCAGTCAGAAGAACgatC-3' |
| Bottom strand | 5'-aaacGatcGTTCTTCTGACTGGCAC-3' |

**Table S8. Top-5 predicted off-target sites**

| sgRNA off-target | Sequence | Chromosome | Score |
|---|---|---|---|
| sgRNA4 off-target 1 | TATCATTCTAAACATCACCA | Chr 6 | 1.714 |
| sgRNA4 off-target 2 | CATCCTTTTAGAAATCACCA | Chr 5 | 1.109 |
| sgRNA4 off-target 3 | CTTCTTTGTAGACAGCACCA | Chr 6 | 0.937 |
| sgRNA4 off-target 4 | CAACTTTAAAGACATCACCA | Chr 8 | 0.883 |
| sgRNA4 off-target 5 | AATCCTTGTGCACATCACCA | Chr 12 | 0.846 |

**Table S9. Sequences of primers to amplify the genomic region of each off-target site**

| PCR Target | Sequence |
|---|---|
| sgRNA4 off-target 1 | Forward primer: 5'-GACCCGTAGCAAAGCCAAG-3' |
| | Reverse primer: 5'-AATAGAAATGTCAGACCCAGTGA-3' |
| sgRNA4 off-target 2 | Forward primer: 5'-CCAGTCCCTGTGTGTTTTCG-3' |
| | Reverse primer: 5'-TCATCCTCTATCCCGGTAGTT-3' |
| sgRNA4 off-target 3 | Forward primer: 5'-ACCCTGGTTTTGAGTACTCCT-3' |
| | Reverse primer: 5'-AGCAGCCCTCCTAACTTTCT-3' |
| sgRNA4 off-target 4 | Forward primer: 5'-TCCCTCTATGTTCACCAGCA-3' |
| | Reverse primer: 5'-TCCCGTGCTACAATTCTTTATTTAA-3' |
| sgRNA4 off-target 5 | Forward primer: 5'-TTTCAGCTTTCCCTGCATTT-3' |
| | Reverse primer: 5'-CACTCTGAGCCCATTTCACC-3' |
